# RNA G-quadruplex folding is a multi-pathway process driven by conformational entropy

**DOI:** 10.1101/2023.02.08.527282

**Authors:** Marijana Ugrina, Ines Burkhart, Diana Müller, Harald Schwalbe, Nadine Schwierz

## Abstract

The kinetics of folding is crucial for the function of many regulatory RNAs including RNA G-quadruplexes (rG4s). Here, we characterize the folding pathways of a G-quadruplex from the telomeric repeat-containing RNA by combining all-atom molecular dynamics and coarse-grained simulations with circular dichroism experiments. The quadruplex fold is stabilized by cations and thus, the ion atmosphere forming a double layer surrounding the highly charged quadruplex guides the folding process. To capture the ionic double layer in implicit solvent coarse-grained simulations correctly, we develop a matching procedure based on all-atom simulations in explicit water. The procedure yields quantitative agreement between simulations and experiments as judged by the populations of folded and unfolded states at different salt concentrations and temperatures. Subsequently, we show that coarse-grained simulations with a resolution of three interaction sites per nucleotide are well suited to resolve the folding pathways and their intermediate states. The results reveal that the folding progresses from unpaired chain via hairpin, triplex and double-hairpin constellations to the final folded structure. The two- and three-strand intermediates are stabilized by transient Hoogsteen interactions. Each pathway passes through two on-pathway intermediates.

We hypothesize that conformational entropy is a hallmark of rG4 folding. Conformational entropy leads to the observed branched multi-pathway folding process for TERRA25. We corroborate this hypothesis by presenting the free energy landscapes and folding pathways of four rG4 systems with varying loop length.

## INTRODUCTION

RNA G-quadruplexes (rG4s) play a crucial role in a variety of physiological processes including regulation of transcription and translation (1, 2) or pre-mRNA processing (3, 4). G4-forming sequences are abundant in the human genome and non-canonical structures of guanosine-rich sequences can be formed by DNA and RNA. While long debated (5), advances in the development of highly sensitive approaches provided further evidence for the existence of rG4s in vivo and revealed their dynamic nature in cells (6). However, to resolve how rG4 folding and unfolding affects its function, a detailed understanding at the molecular level is required.

The overall architecture of G4s is striking: Four non-neighboring guanosine nucleobases assemble in quartets and several quartets stack onto one another to form a central channel. For rG4s, this channel is typically flanked by propeller loops, while a large variety of different loop arrangements have been reported for DNA G-quadruplexes (dG4s) (7). The three-dimensional structures are stabilized by cations. In particular, K^+^ and Na^+^ are frequently observed in the channel and lead to the highest stabilization efficiency against denaturation among alkali and alkaline earth cations (8).

The telomeric repeat-containing RNA (TERRA) is an important rG4 example. Telomeres are specialized nucleoprotein structures that protect chromosomal DNA from progressive degradation. Transcription of the telomeric C-rich strand in the chromosomes produces TERRA, which has a canonical G-rich motif with sequence r(UUAGGG) (9, 10). TERRA is essential in maintaining genomic integrity by regulating telomerase activity and protecting the chromosome ends against degradation (11, 12). Telomere dysfunction is connected to cell aging and cancer (13, 14) and the design of small drug molecules targeting TERRA attracts therefore increasing scientific attention (15).

Despite the importance of rG4s, most scientific work has focused on dG4s so far (16). The scientific focus on dG4s can be rationalized by the fact that their existence in vivo has been demonstrated early on by different experimental techniques (17, 18). By contrast, the existence of rG4s in vivo and their biological relevance are just now starting to be established (2, 6, 19, 20). In addition, the investigation of rG4s requires much more sensitive experimental techniques since rG4s are more dynamic compared to their DNA counterparts (6).

Moreover, the folding intermediates of rG4s are transient, making their detection and resolution a major challenge (21). Consequently, the molecular details of folding have not been resolved so far and the intermediate states remain elusive.

Insights into the atomistic structure of folded G4s are obtained from nuclear magnetic resonance (NMR) spectroscopy (22–25) and X-ray crystallography (26). In addition, time-resolved NMR spectroscopy yields information on possible folding intermediates (27–29). Such experiments recently revealed that the folding kinetics of TERRA is fundamentally different compared to a homologous DNA sequence (30, 31). The K^+^-induced folding kinetics of TERRA is an order of magnitude faster (k_1_ = 1.45*/*min) compared to the dG4 of identical nucleotide sequence (k_1_ = 0.41*/*min) (21).

While some dG4 intermediates have been resolved at atomistic resolution by NMR (23, 32–34), the intermediate states of rG4s remain elusive due to their short lifetimes (27). Further information on folding intermediates can be obtained from single-molecule techniques such as single-molecule Förster resonance energy transfer (smFRET). Based on the measured distance between strategically placed fluorophores, hairpin and triplex structures have been suggested as intermediates of two and three quartet rG4s and dG4s (35, 36).

To complement experiments, biomolecular simulations are powerful tools which provide molecular insights into the folding pathways and intermediate states of rG4s and dG4s (16, 37–41). At first sight, all-atom molecular dynamics (MD) simulations in explicit water seem to be the ideal computational method. For instance, atomistic simulations allow to resolve the conformational rearrangements in the folded state of G4s, their interactions with different cations, (42–44) or the stability of putative intermediates along the folding pathway (37, 38). Moreover, combining atomistic simulations and enhanced sampling techniques such as replica exchange can improve the sampling of rare folding events (37–40). Still, G4 folding is on the timescale of minutes. Simulating a single folding pathway, let alone the large variety of different pathways, is therefore out of reach for atomistic simulations.

The limitations of atomistic simulations can be overcome by coarse-grained (CG) simulations. CG models simplify the atomistic representation by reducing the number of particles to three to seven beads per nucleotide and employing implicit solvent thereby reducing the computational costs further. For instance, the HiRE-DNA model revealed the potential of CG model to investigate dG4 folding by resolving numerous intermediates of human telomeric dG4s (41). By now a variety of CG models for RNA are available such as SimRNA (45), HiRE-RNA (46), seRNA (47) and TIS model (48–53). TIS, the three-interaction site model developed by Thirumalai and coworkers, is a simple low-resolution model. Its efficiency and accuracy to describe folding has been demonstrated for a variety of RNA molecules ranging in size from larger molecules such as ribozyme (52) to smaller RNAs like riboswitch (54), pseudoknot (50), and hairpin (49).

By combining CG simulations using the TIS model and circular dichroism (CD) experiments, we resolve the folding pathways and intermediate states of the TERRA rG4 sequence consisting of a monomeric 25-mer human telomeric RNA repeat. In a first step, we validate the CG model by CD experiments and obtain close agreement as judged by the population of folded states as a function of temperature and salt concentration. Quantitative agreement is obtained by a newly developed concentration matching procedure, which takes the ionic double layer into account via additional all-atom MD simulations. Subsequently, we resolve the folding pathways from the CG simulations and identify hairpins, triplexes and double-hairpins as intermediate states. Finally, we investigate the folding of three synthetic rG4 sequences with varying loop length and a plant rG4 sequence and confirm that branched multi-pathway folding is a characteristic feature of all rG4 systems.

## METHODS

### Coarse-grained and atomistic simulations Simulation models

We modeled the monomeric 25-mer G4 from human telomeric RNA (TERRA25). As starting point, we used the NMR resolved structure of a dimeric 24-mer TERRA G4 in K^+^ solution (PDB-ID: 2KBP from ref. (55)) with sequence (UAGGGUUAGGGU)_2_. In a first step, we used the ModeRNA homology modeling server (56) to connect the two strands of the RNA into a single strand through U12 and U13. We created an additional propeller loop and added a U25 to the 3’-end of the strand. In a second step, we optimized the structure using the SimRNA server (45) and selected 10 minimum energy structures. We preformed MD simulations of the 10 structures for 100 ns and selected the one with lowest root mean square deviation (RMSD) with respect to the experimental structure (2KBP).

The equilibrated TERRA25 structure with sequence 5’-UAGGGUUAGGGUUAGGGUUAGGGUU-3’ is shown in Figure 1A. It consists of four guanosine repeats. Each repeat is formed by three consecutive guanosines in anti-conformation (i.e. with a glycosidic torsion angle between 180° and 240°). The repeats are connected by propeller loops. The guanosines from the four repeats form 3 quartets that give the G4 its characteristic non-canonical structure.

**Figure 1.**
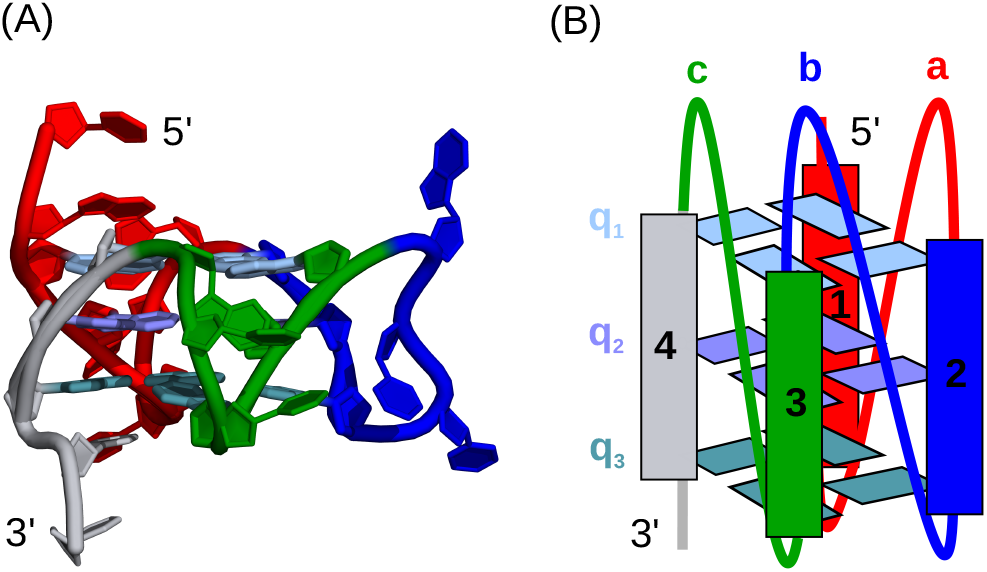
Monomeric 25-mer TERRA G4 with sequence 5’-UAGGGUUAGGGUUAGGGUUAGGGUU-3’ in the simulations. Simulation snapshot from all-atom MD simulations (A) and schematic representation (B). Each of the four GGG-repeats is represented in a different color (from 5’ to 3’: red (1), blue (2), green (3), gray (4). The three G-quartets consist of four guanosines from different repeats and are labeled q_1_, q_2_, q_3_ in (B). The three propeller loops are labeled a, b, c. The connecting loops all have the sequence UUA. The sequence has a UU-overhang at the 3’ end and a UA-overhang at the 5’ end.

To test if the branched multi-pathway folding process observed for TERRA25 also occurs in different systems, we investigated four additional rG4 systems (Table 1). The first three systems are synthetic sequences of varying loop lengths. They contained only uridine nucleosites in the loops 5’-UAG_3_U_n_G_3_U_n_G_3_U_n_G_3_UU-3’, where *n* = 1, 2, 3. The fourth system is a plant rG4 sequence that was previously studied by smFRET (36) and has the sequence 5’-GGGAGGGAAGGGGAAGGGG-3’. We employed the ModeRNA server (56) to create the initial folded structure and used the structure of TERRA25 as a reference. Since the last structure has more than 3 guanosines in the last two repeats, we used RNAFold web server from the ViennaRNA Package 2 (57) to model the secondary structure and generated the native structure accordingly.

**Table 1.**
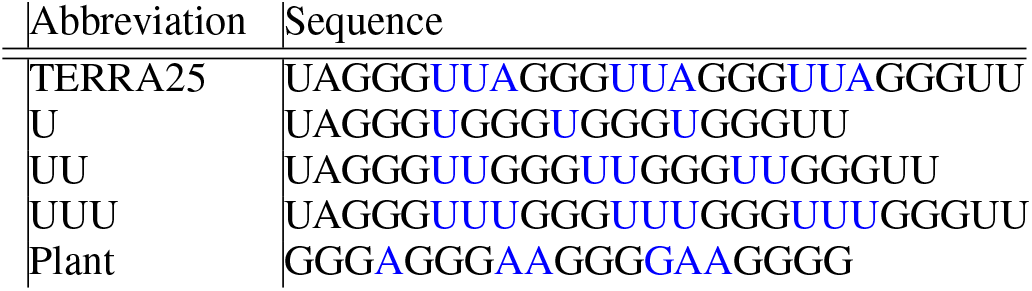
Sequences of rG4 structures simulated in this work. Loop regions are marked in blue.

### All-atom MD simulations

The MD simulations were performed with Gromacs version 2018.1 (58). Periodic boundary conditions were used with the particle mesh Ewald method (59) to calculate long range electrostatics, with cubic interpolation and a Fourier spacing of 0.12 nm. The cutoff for Coulomb and Lennard-Jones interactions was 1.2 nm and long-range dispersion corrections for energy and pressure were applied. Bond to hydrogens were constrained with the LINCS algorithm (60) with order of four for the matrix inversion correction and one iterative correction. Simulations were performed for 100 ns with a 2-fs time step. TERRA25 was simulated using parameters from Amber99sb -ildn* force field (61) with parambsc0 (62) and *χ*_OL3_ (63) corrections and TIP3P water model (64). Optimized parameters were used for K^+^ and Cl^−^ ions, which correctly reproduce the thermodynamic and kinetic properties of the ions in TIP3P water (65). We added the same number of anions and cations to obtain different bulk concentrations and 24 additional cations were used to neutralize the systems.

In the first set of simulations, we simulated the 10 conformations obtained from simRNA for which the interatomic potential energy is lowest. 1 M KCl was used in each of the 10 systems. The structures were initially placed in a rectangular box with dimensions 5.4*×* 5.0*×* 5.6 nm^3^. From these simulations, the equilibrated structure with the lowest RMSD with respect to the experimental structure was selected for all further simulations. In the RMSD calculations only heavy atoms from the central pore were used.

In a second set of simulations, we calculated the ion distribution profiles for TERRA25 to establish the relation between bulk salt concentration and the local concentration in the double layer.

We performed simulations of TERRA25 at eight concentrations of KCl, namely 0.48 mM, 16 mM, 53 mM, 82 mM, 0.14 M, 0.22 M, 1.03 M, and 1.89 M. For the 0.48 mM KCl simulation, the box size with edge length 30 nm was used. It contained 878.660 water molecules. For the 16 mM KCl simulation, a box size with edge length 15 nm was used. The box contained 108.299 water molecules. For the KCl concentrations 0.22 M and 1.89 M a box size with edge length of 10.0 nm was used. These systems contained 32.108 or 30.116 water molecules. For the 53 mM, 82 mM, 0.14 M, and 1.03 M KCl concentrations, we used a smaller box with edge length 9.0 nm. The boxes contained between 22.744 and 23.551 water molecules.

To investigate if the matching procedure for KCl is also valid for NaCl, we performed one additional 100 ns simulation of TERRA25 in 1 M NaCl using the Mamatkulov-Schwierz parameters (65). The simulation box had an edge length of 9 nm, and it contained 22.744 water molecules. RMSD, simulation snapshots, ion distribution profiles are shown in the Supporting Information (Figure S1 and S2).

TERRA25 was placed in a cubic box and each system was equilibrated in NVT and NPT simulations. During equilibration harmonic restraints with force constant 1000 kJ/(mol nm^2^) were applied on the heavy atoms of TERRA25 to prevent large conformational changes. The NVT equilibration was done in two parts. During the first 1 ns restraints of 2000 kJ/(mol nm^2^) were applied to ions to prevent ion pairing before a complete hydration shell is formed. In the subsequent 1 ns, the restraints on the ions were released. As thermostat the velocity rescaling algorithm with a stochastic term (66) and with a coupling constant of 0.1 ps was used. During the 2 ns NPT equilibration, pressure coupling was performed using the Berendsen barostat (67) with a 1 ps coupling constant. In the production runs, the velocity rescaling thermostat (66) and the Parrinello-Rahman barostat (68) with a 2 ps coupling constant were used. Position restraints on TERRA25 were released during the production run. Each production run was 100 ns long. The radial concentration profile was calculated as follows: First, a 2D number density profile of K^+^ around the central pore of the G4 was calculated using Gromacs and an in-house code (69). The radial concentration profile was obtained by integration and normalization.

### CG simulations using TIS

The CG simulations were performed using the Three Interaction Sites (TIS) model developed by Thirumalai and coworkers (48–53). In the TIS model, each nucleotide is represented by three beads, located at the center of mass of the phosphate group, sugar and nucleobase. The parameters for the canonical interactions between the beads were taken from previous work (48). The intramolecular attractive interactions were defined based on the residues that appear in the native structure of TERRA25. This ensures that the native structure is the minimum energy structure (70). Native hydrogen bonds and tertiary stacks were defined based on the modelled monomeric TERRA25 structure described above.

In order to capture folding intermediates that are not stabilized by native interactions, non-native secondary structure interactions were included via the base-stacking interactions of consecutive nucleotides and hydrogen-bond interactions between all nucleobases as in previous work (53). The parameters for these interactions were calculated by coarse-graining the standard A-form RNA helix (48).

A non-interacting uridine was added capping the structure at the 3’ end. The RNA was placed in a cubic box with an edge length of 70 nm.

The solvent was modeled implicitly, and we employ the Debye-Hückel approximation in combination with the concept of counterion condensation to take the ions into account (48). Specifically, the contribution to the energy function of TIS for the interaction between two phosphate groups is based on the linearized Debye-Hückel equation and is given by (48, 71)

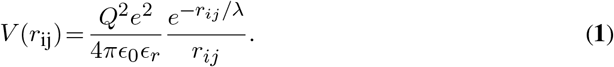

Here, *r*_*ij*_ is the distance between the two phosphate groups *i* and *j, Q* is the charge of a phosphate group, *e* is an elementary charge, *ϵ*_0_ the dielectric constant of vacuum, *ϵ*_*r*_ is the dielectric constant of water and *λ* is the Debye length. Hence, the ionic concentration is included in the CG model via the Debye length

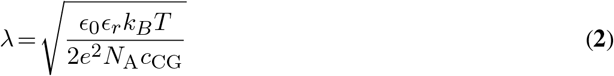

where *N*_A_ is Avogadro’s constant and *c*_CG_ is the uniform ion concentration of the CG model.

Numerical integration of the equations of motion was performed using the leap-frog algorithm with time step h= 0.05*τ*, where *τ* = 50 fs as in previous work with the TIS model (49). The simulations were performed in the low friction regime to increase the sampling efficiency by reducing the viscosity of water by a factor of 100 (72). The cut-off for electrostatic interactions was set to 3 nm.

To investigate the temperature dependence of the population of folded rG4 states, we performed 120 simulations of independent copies of the system at each of temperatures in the temperature range from 1-120°C, leading to a temperature spacing between the copies of 1°C.

Each temperature was simulated at a concentration of *c*_CG_ = 150 mM (corresponding to *c*_bulk_ = 12 mM) for about 100 *μ*s.

To investigate the concentration dependence of folding, we performed independent simulations of copies of the system at 16 different concentrations in a range of from 10 *μ*M to 1 M (corresponding to *c*_bulk_ = 0.06 *μ*M to *c*_bulk_ = 430 mM) at two different temperatures, 25°C and 60°C. Each setup was simulated for about 100 *μ*s. The simulations were initiated from the folded state and 5 *μ*s were neglected for equilibration in the analysis. To provide sufficient statistics at T=25°C, additional simulations were performed at 0.1, 0.4 and 1 mM using 50 additional independent simulations of 20 *μ*s.

The folding and unfolding pathways were simulated at room temperature (T=25°C) and at *c*_CG_ = 50 mM (corresponding to *c*_bulk_ = 0.3 mM). The salt concentration was chosen such that the probabilities of folded and unfolded structures are similar. The plant rG4 was simulated at *c*_CG_ = 400 mM (corresponding to *c*_bulk_ = 80 mM) and 25°C. In order to provide sufficient statistics, 100 copies of system were simulated. For 23 copies of the system, we performed 180 *μ*s long simulations and for 77 copies we performed 52.5 *μ*s long simulations. In total, 8182.5 *μ*s of simulation time was used.

The simulations were initiated from the folded state and 5 *μ*s were neglected for equilibration in the analysis. We observed about two folding/unfolding events per trajectory and the number of forward and backward transitions during 180 *μ*s simulations were the same.

### Intermediate states in TIS

A common simplification used in Gō-like models is that intramolecular attractive interactions are defined only between the residues that appear to be in contact in the native structure. This definition ensures that the native structure is the minimum energy structure. The drawback of such basic Gō-like models is that they cannot capture partially folded intermediate states stabilized by non-native interactions. The TIS model, used in this current work, is more advanced than standard Gō-like models. TIS explicitly includes non-native secondary structure interactions (48). In particular, all base-stacking interactions between consecutive nucleotides and hydrogen-bond interactions between any bases G and C, A and U, or G and U are included (48). Therefore, the model captures non-native secondary structure interactions as shown in previous work (54, 73) and for TERRA25 (see Figure S3). Moreover, it has been shown that TIS reproduces experimental thermodynamic and structural data for several different RNA molecules including ribozyme formation or riboswitch folding under a variety of solvent conditions (52, 54, 73). With these prerequisites, TIS is ideal to resolve intermediates states and their frequency of occurrence in the folding of rG4 systems.

### Timescale, rates, and lifetimes in TIS

The dynamics of the system is based on the Langevin equation. The effect of the solvent is modeled implicitly via a Gaussian random force and a friction force which is assumed to be proportional to the viscosity *η* of the medium. The Langevin equation is solved numerically using a finite timestep ∆*t* = 2.5 fs. The dynamics obtained from the solution of the Langevin equation sets the timescale of the coarse-grained simulations. However, relating the timescale of CG models to the timescale observed in experiments is challenging for two reasons: (i) The viscosity of the water in the CG simulations is reduced by a factor of 100 to enhance the conformational sampling as in previous work (48). (ii) The free energy folding landscape is smooth due to the coarsening of atomistic interactions and the implicit solvent.

One possibility to relate the scales is to introduce a time rescaling factor based on experimental folding rate (21)

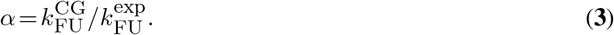

With 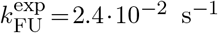 and 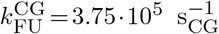, we obtain *α* = 1.56·10^7^ for TERRA25. The corresponding rates and lifetimes are listed in Table S2 and S3 in the Supporting Information.

However, a single numerical factor is unlikely to capture the coarsening of complex atomistic interactions. We therefore provide the results in CG time units *s*_CG_ and introduce reduced rates 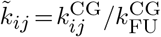and lifetimes 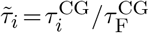in addition to the rescaled results (Table S2 and S3).

The kinetic rate coefficients *k*_ij_ for transitions *j*→ *i* are calculated from

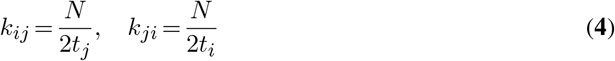

where *N* is the total number of transitions between the two stable states and *t*_j/i_ is the total time in the respective state. Note that for long equilibrium runs, the number of forward and reverse transitions are equal. Further information and the rate equations for the time evolution of the individual states can be found in the Supporting Information (section Rate equations for rG4 folding).

For first-order kinetics, the lifetime distribution is determined by the rates. Specifically, the mean lifetime of state *j* is calculated from

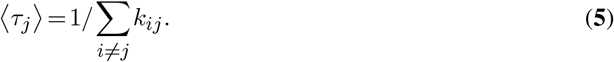

### Gibbs free energy

To determine the stability of the different rG4 systems the Gibbs free energy was calculated via

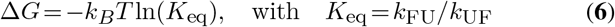

where K_eq_ is the equilibrium constant. The results are compared to the results from UV melting experiments (74). Since our simulations were performed at 298.15 K, we calculate the Gibbs free energy 298.15 K from the experimental entropy change and experimental enthalpy change (see Supporting Information, section Gibbs free energy from CG simulations and thermal melding curves for details). Moreover, the experimental results are the average over different loop sequences and the concentration of RNA and KCl ions were different in experiments and simulations. In the following, we therefore calculate the Gibbs free energy change with respect to a reference system. For the simulations, we chose the UUU system and the L333 system ref. (74) for the experiments.

### Number of native contacts and radius of gyration

To identify the transition between the folded and unfolded state, we calculated the radius of gyration *r*_*g*_ and the number of native contacts *n*_*C*_ between the guanosines in each quartet. *n*_*C*_ = 12 corresponds to the fully folded, native state. *n*_*C*_ is calculated using the coordination function of PLUMED (75) with cutoff distance between guanosine base beads *r*_0_ = 0.75 nm, offset parameter *d*_0_ = 0.08 nm and parameters *n* = 35 and *m* = 55.

### Order parameters

To characterize the folding intermediates, we introduce two order parameters. The repeat order parameter counts the number of native contacts between repeats *i* and *j* (with *i*≠ *j* and *i,j* = 1−4). It is calculated from the indicator function

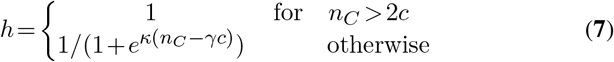

where *n*_*C*_ is the number of native contacts between repeats *i* and *j* and the parameters *κ* = −5, *γ* = 1.3, *c* = 1.4. This choice provides a smooth function for the order parameter without fluctuations. The repeat order parameter *r*_*ij*_ = *h* is one if all three native contacts have formed between the neighboring repeats *i* and *j*.

To resolve the twelve different intermediate states, we introduce two additional order parameters, *s*_1_ and *s*_2_, based on linear combinations of *r*_*ij*_

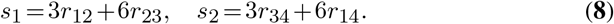

The values for the repeat order parameters and *s*_1_ and *s*_2_ are illustrated Figure S4. The quartet order parameter *q*_*k*_ counts the number of native contacts in each of the three quartets (*k* =−1 3). *q*_*k*_ is calculated from equation **7** with *κ* =− 4, *γ* = 1.9, *c* = 1.93, and *n*_*C*_ being the number of native contacts in quartet *k*. The quartet order parameter is one if all four native contacts have formed in a quartet.

### Fraction of folded structures

For the fraction of folded structures *n*_*f*_, we calculated the total number of contacts between the guanosines and use equation **7** with *κ* = −5, *γ* = 2 and *c* = 4.5. *n*_*f*_ is one in the folded state. The fraction of folded structures was calculated from the time the molecule spends in the folded state relative to the total time. The error is calculated by block averaging over 2.5 *μ*s blocks.

### Concentration matching

In the CG simulations, the solvent is modeled implicitly, and the concentration of ions is homogeneous in the simulation box (Figure 2B). Due to the highly charged backbone of the rG4, cations are attracted to the phosphodiester backbone and form a pronounced electric double layer (Figure 2A). Since screening of ions in the vicinity of phosphodiester groups of RNA is a driving factor in RNA folding, the concentration in the CG model needs to be matched to the local concentration of ions in the electric double layer.

**Figure 2.**
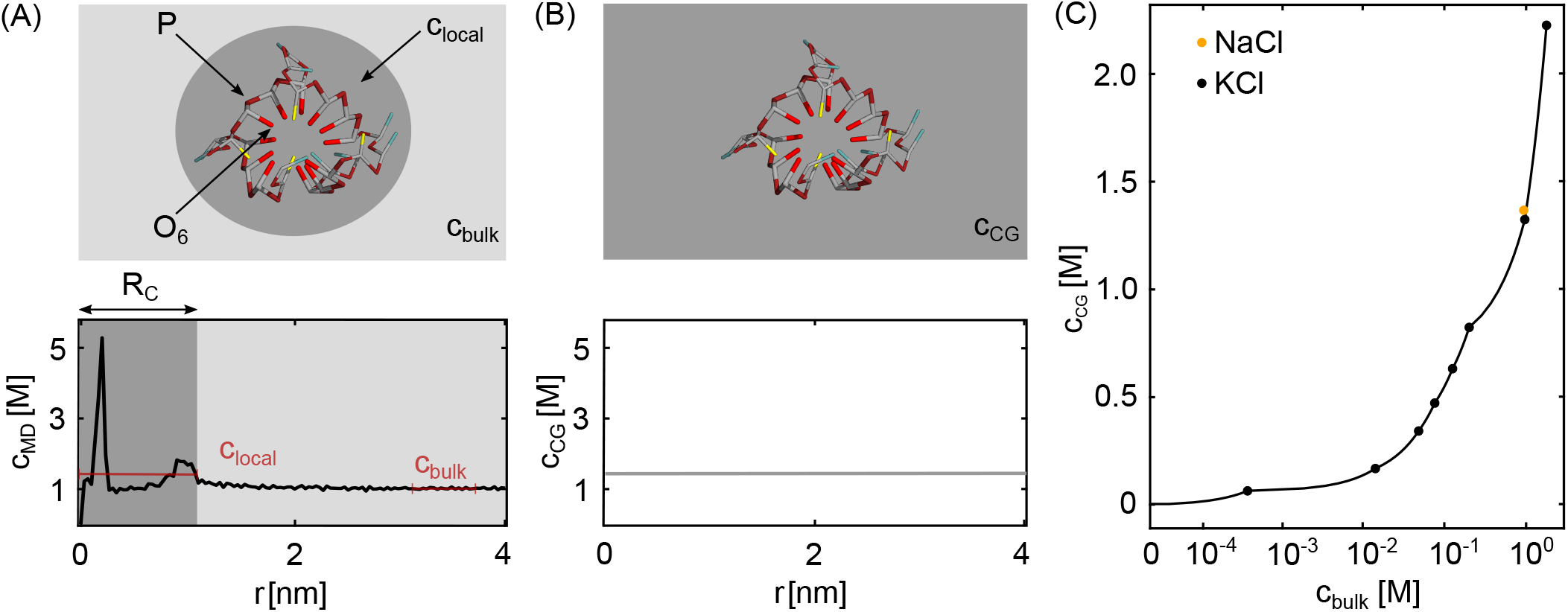
Concentration matching to mimic the effect of an ionic double layer in implicit solvent CG simulations. (A) Schematic representation of the ion atmosphere around the rG4 (top) and concentration profile *c*_MD_ (*r*) of K^+^ ions as function of the distance from the central channel from all-atom MD simulations (bottom). *c*_bulk_ is the bulk salt concentration and *c*local the averaged concentration in the double layer within *R*_C_. (B) Uniform concentration profile of the CG model *c*_CG_ = *c*_local_ after matching to the local concentration in the double layer obtained from the MD simulations. (C) Matching relation of the bulk concentration *c*_bulk_ and the CG concentration *c*_CG_. The matching is done for KCl. One additional all-atom MD simulation was performed for *c*_bulk_ = 1 M NaCl.

To achieve this, we developed the following matching procedure: Firstly, we derive the correlation between the bulk concentration (as measured in the experiments) and the local concentration in the double layer by performing all-atom MD simulations. The simulations were done at eight different KCl concentrations. Figure 2A shows the radial concentration profile at *c*_bulk_ = 1.03 M. The formation of an electric double layer is clearly visible. The highest peak (*r* = 0.2 nm) corresponds to K^+^ in the central pore and the second peak (*r* = 1.23 nm) corresponds to adsorption of K^+^ at the phosphate oxygens of the backbone. From the simulated profiles, we calculated the local concentration by averaging from the center of the channel up to the distance *R*_C_. The distance *R*_C_ is a free parameter representing the distance of the edge of the bilayer from the geometrical center of the RNA. Since the results are not sensitive to the exact choice of *R*_*C*_, we chose *R*_C_ = 1.5 nm, which best reproduced the experimental results from Figure 3. We calculated the bulk concentration by averaging the concentration profiles over the last 0.5 nm from the edge of the simulation box. Due to the high binding affinity of K^+^, the bulk concentration vanishes in finite simulations boxes at low concentrations. To provide consistent results, we therefore used the anion profiles to determine the bulk concentration (Figure S2).

**Figure 3.**
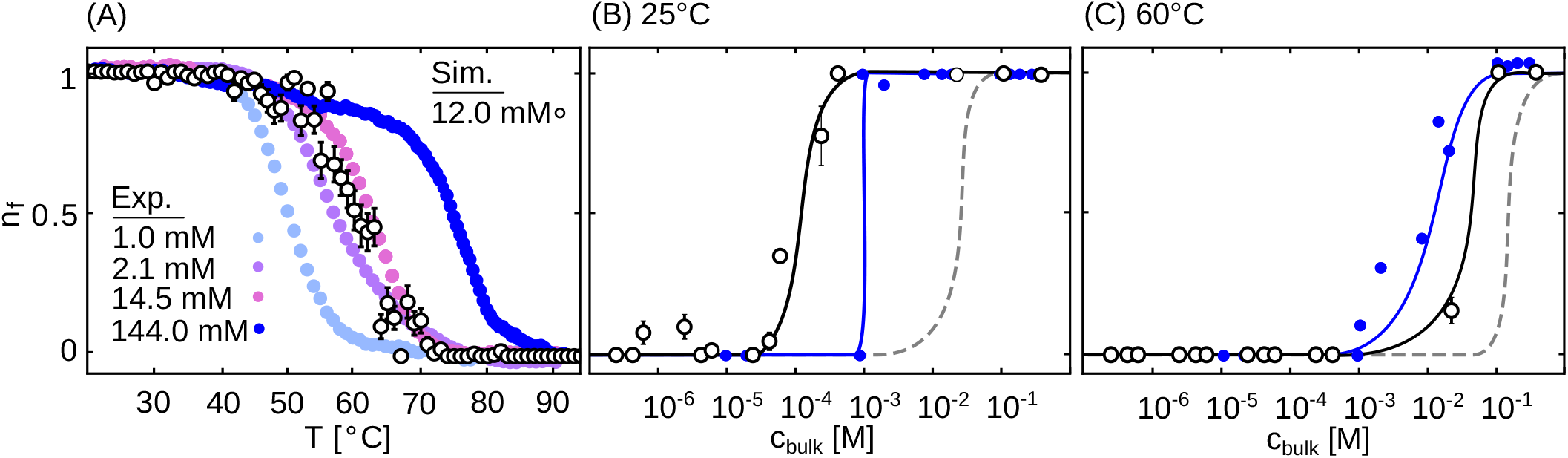
Comparison of the population of the folded state *n*_f_ from simulations and experiments. (A) *n*_f_ as function of the temperature from experimental melting curves and CG simulations. (B, C) *n*_f_ as function of the K^+^ concentrations at 25°C and 60°C. With increasing salt concentration, the equilibrium is shifted to the folded state with *n*_f_ = 1. The lines are a guide to the eye and indicate the folded/unfolded transition region. The dashed lines are the uncorrected results neglecting the effect of the ionic double layer.

We performed a linear interpolation between the datapoints, and the resulting matching relation is shown in Figure 2C and Table S1. The results from the matching procedure are validated by CD experiments as will be discussed further below.

### Experiments

#### Sample preparation

TERRA25 (5’-UAGGGUUAGGGUUAGGGUUAGGGUU-3’) was purchased from Dharmacon (Cambridge, UK). The oligomer was HPLC purified using a tetrabutylammonium acetate buffer and precipitated with five volumes of LiClO_4_ (2% in acetone (w/v)) at -20 °C over night. Subsequently, desalting with an ultracentrifuge filtration device (Vivacon 2 kD cut-off, VWR) was performed to remove residual Li^+^ cations and acetone.

#### Circular dichroism (CD) experiments

CD measurements were performed using a JASCO J-810 spectropolarimeter with a Peltier temperature control system. 10 *μ*M RNA were used per measurement and dissolved in ddH_2_O containing 1, 2.1, 14.5 or 144 mM KCl. A quartz cuvette with 2 mm pathlength was used. The CD spectra were recorded with a scanning speed of 50 nm/min and three accumulations between 200 and 320 nm. CD melting curves were recorded at the maximum of the CD signal (264 nm) with a heating rate of 0.5°C/min.

#### Native polyacrylamide gel electrophoreses (PAGE)

Correct folding of TERRA25 was checked via native PAGE (15% acrylamide gel). 400 pmol samples were loaded in 50% glycerol and 5 mM KCl. Gels were prepared in 50 mM Tris-borate buffer (pH = 8.4) supplemented with 5 mM KCl. Bands were separated in the same buffer at 0.8 W for 90 min with water cooling (13°C water temperature). The bands have been visualized by Stains-All.

#### Fraction of folded structures from CD melting curves

Conversion of CD absorbance into population of the folded state was calculated according to the method highlighted by Mergny and Lacroix (76). Firstly, upper and lower baselines were corrected for temperature correction to determine the CD signal for the unfolded and folded states, respectively. The baselines were then used to normalize the absorbance values and yield fraction of folded states.

The melting temperatures were calculated by fitting the normalized melting curves to a sigmoidal function

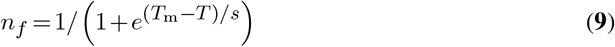

where *T*_m_ is the melting temperature and *s* the slope of the curve. The fitting was done according to the nonlinear least-squares Marquardt-Levenberg algorithm.

## RESULTS AND DISCUSSION

### Quantitative comparison of the fraction of folded structures from simulations and experiments

Initially, we validated the CG model by CD experiments and provided a direct comparison of the fraction of folded structures in dependence of the temperature or the K^+^ concentration (Figure 3). To obtain quantitative agreement between experiments and CG simulations, the ion atmosphere surrounding the highly charged G4 has to be taken into account. Since the cations are attracted to the negatively charged phosphodiester backbone while the anions are repelled, a pronounced double layer is formed. Given the high negative charge, the local ion concentration in the double layer is significantly larger compared to bulk (Figure 2C). It is therefore crucial to take the local concentration in the CG simulations into account to correctly capture the population of the folded state at a given bulk salt concentration. To achieve this, we derived a matching relation from all-atom MD simulations which relates the bulk salt concentration to the local concentration in the double layer (Figure 2C). Based on the relation, the CG concentration was chosen such that it reproduces the average concentration in the double layer.

To validate the procedure, we first investigated the temperature-induced unfolding of the TERRA25 and obtained melting curves from experiments and simulations (Figure 3A). At low temperatures, rG4 is folded in presence of K^+^ (*n*_*f*_ = 1). With increasing temperature, the equilibrium shifts toward the unfolded state and *n*_*f*_ decreases. At high temperatures, rG4 is unfolded (*n*_*f*_ = 0). The melting point *T*_*m*_ depends on the salt concentration (Table 2) and the rG4 becomes more stable with increasing K^+^ concentration. The direct comparison of experiments and simulations reveals a similar dependence of *n*_*f*_ on the temperature (Figure 3A). Moreover, the melting temperature of 60°C from the CG simulations at 12 mM (derived from the matching relation in Figure 2C) is in between the experimental results of 59°C and 62°C for 2.1 and 14.5 mM (see Table 2). Note that without the matching relation, a melting temperature of 60°C corresponds to 150 mM salt concentration. Hence, without the details of the ionic double layer, the CG model clearly overrates the population of the unfolded state.

**Table 2.** Melting temperatures of TERRA25 at different KCl concentrations from CD experiments and CG simulations using concentration matching from all-atom MD simulations.

| | $c_{\text{bulk}}$ [mM] | $c_{\text{CG}}$ [mM] | $T_m$ [°C] |
| --- | --- | --- | --- |
| Exp. | 1.0 | | $50.67 \pm 0.04$ |
| | 2.1 | | $59.21 \pm 0.09$ |
| | 14.5 | | $62.6 \pm 0.2$ |
| | 144.0 | | $76.9 \pm 0.1$ |
| Sim. | 12 | 150 | $60.1 \pm 0.1$ |

The importance of including the local salt concentration via the matching relation becomes even more evident from the dependence of *n*_*f*_ on the concentration at 25°C or 60°C (Figure 3B,C). Without matching, the CG model overrates the destabilization of the folded state, and the folded/unfolded transition is predicted at too high salt concentrations. On the other hand, quantitative agreement of CD experiments and CG simulations is obtained by taking the local concentration in the double layer into account via the matching relation from additional MD simulations.

### Folding and unfolding of TERRA25 is a sequential but multi-pathway process

After successful validation, we analyzed the CG simulations to extract information regarding the folding and unfolding of TERRA25.

Figure 4B shows the number of native contacts *n*_*c*_ and the radius of gyration *r*_*g*_ for an 80 μs simulation at 25 °C and *c*_bulk_ = 0.3 mM. Multiple transitions between the folded (*n*_*c*_ = 12 and *r*_*g*_≈ 15 Å) and unfolded (*n*_*c*_ = 0 and *r*_*g*_ ≈24) states are observed. Energy and entropy contributions to the folding are obtained by calculating the free energy profile *F* (*r*_*g*_) (Figure 4C). The folded state is characterized by the sharp and deep minimum in the free energy profile while the unfolded state is broader with *r*_*g*_ ranging from 15 to 25 Å . This reflects an energy-entropy compensation mechanism in which the folded state is energetically favored while the unfolded state competes with a larger entropy due to a variety of unfolded conformations. The folded and unfolded structures are separated by a barrier of 5 k_B_T. The two-dimensional free energy landscape *F* (*n*_*c*_,*r*_*g*_) in the region of this barrier reveals several distinct intermediate states (Figure 4C, inset). The intermediate states are broad indicating that multiple different conformations contribute to them (as will be discussed in detail further below).

**Figure 4.**
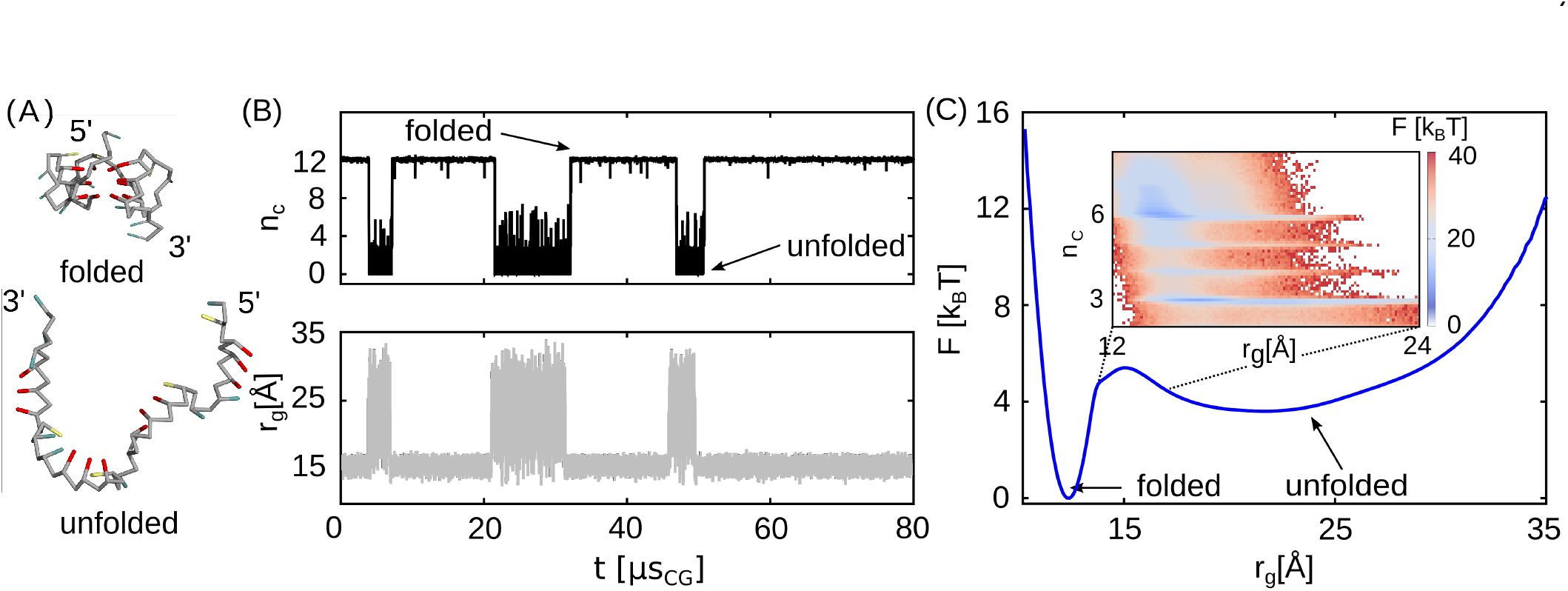
Folding pathway and folding free energy landscape of TERRA25 from CG simulations. (A) Snapshots in the folded and unfolded state from the CG simulations. Guanosine is represented in red, uridine in teal and adenosine in yellow. (B) Number of native contacts *n*C and radius of gyration *r*_g_ as function of simulation time. (C) One-dimensional free energy profile *F* as a function of *r*_g_ . The inset shows the two-dimensional free energy landscape as function of *r*_g_ and *n*C in the region of the barrier.

Figure 5 shows the computation of one representative folding pathway of TERRA25. The number of native contacts *n*_*C*_ increases stepwise while the radius of gyration *r*_*g*_ decreases (Figure 5A). The repeat order parameter *r*_*ij*_ and the quartet order parameter *q*_*i*_ reveal that rG4 is formed repeat after repeat and not quartet after quartet (Figure 5B,C), suggesting that stacking interactions drive the initial formation of rG4 rather than Hoogsten interactions. In this example, repeat 1 and 4 assemble first and form the parallel hairpin H4 without propeller loop (Figure 5D,E at t=47 ns_CG_). Subsequently, repeat 3 assembles and forms the triplex structure T3 with three planar units formed by three h-bonded guanosines, called triads. At this point, the propeller loop c is formed at the 3’ end (Figure 5D,E at t=125 ns_CG_). Folding is completed once the last repeat assembles, creating the loops a and b (Figure 5D,E at t=145 ns_CG_) and all three quartets are formed simultaneously as visible from the quartet order parameter in Figure 5C.

**Figure 5.**
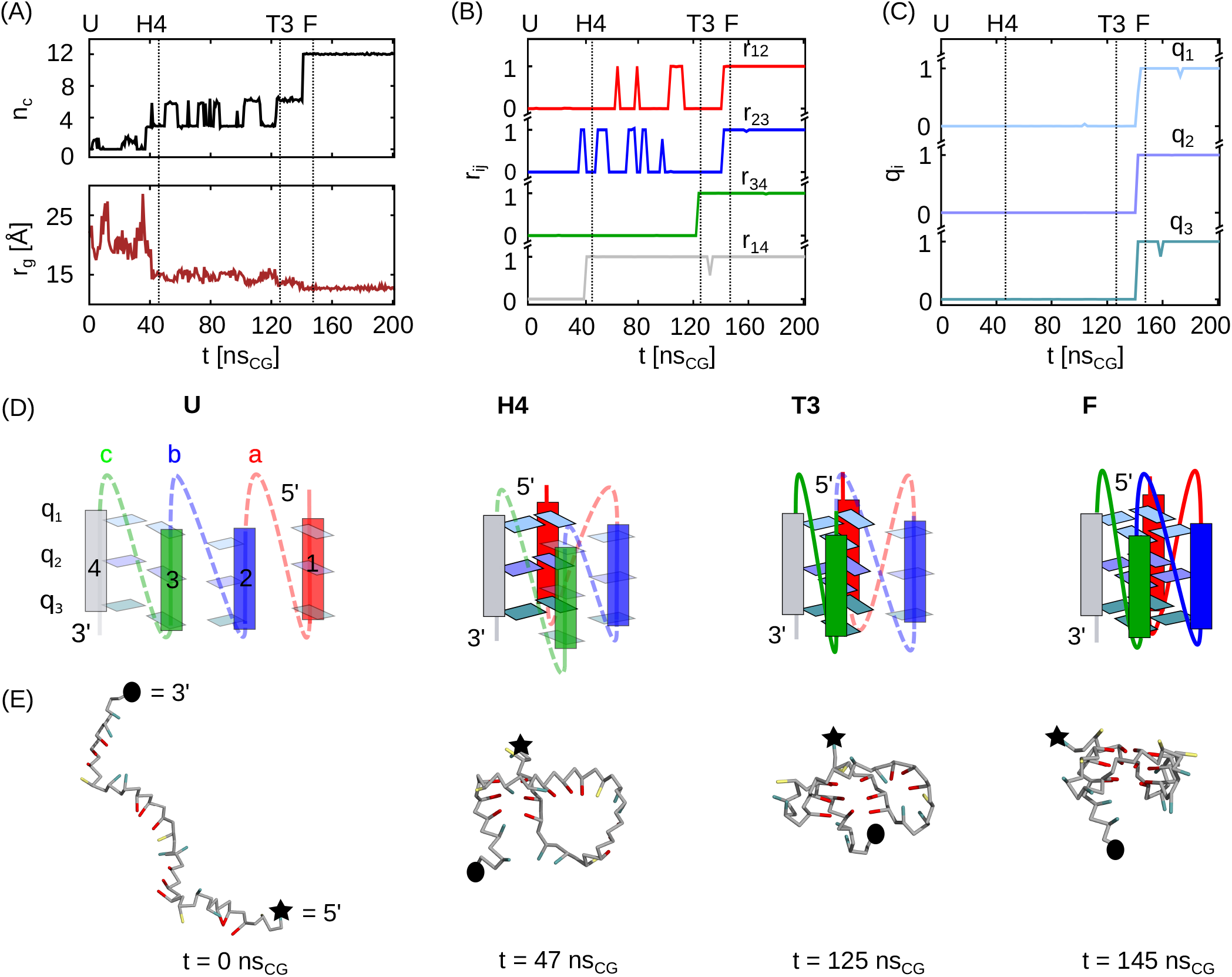
Representative folding pathway and intermediates of TERRA25 from CG simulations. (A) Number of native contacts *n*_C_ and radius of gyration *r*_g_ during the folding transition. (B) Repeat order parameter r_*ij*_ describing the assembly of repeat *i* and *j* (with *i* ≠*j* and *i,j* = 1− 4). r_*ij*_ is one if three native contacts have formed neighboring repeats. (C) Quartet order parameter *qi* describing the formation of the guanosine quartets (*i* = 1 −3). *q*_*i*_ is one if all four native contacts have formed in a quartet. (D) Schematic representation of the folding pathway. Assembled parts of the structure (repeats, quartets, propeller loops) are shown in solid colors; non-assembled parts are transparent or dashed. (E) Simulation snapshots along the folding pathway at times indicated in A-C.

In summary, during rG4 folding the repeats assemble and the corresponding propeller loops are formed. During the folding, two intermediate structures are observed, a hairpin and a triplex, supporting previous observations and predictions of hairpin and triplex structures as intermediates in the folding of rG4s and dG4s in experimental (27, 29, 35, 77) and computational (39, 78) work. In addition, we observe intermediates with non-native interactions (Figure S3). These off-pathway intermediates are transient and unfold quickly and were therefore not analyzed in more detail.

From simple combinatorics, there are four possibilities to form a hairpin from a nucleic acid structure with four repeats since there are only parallel orientations in rG4s (16, 21). Similarly, there are four possibilities to form a triplex. For the folding of G4s the question arises whether all these intermediates can be observed or whether their formation is inhibited kinetically or energetically.

To gain further insights, we determined the folding pathways from CG simulations and classified the intermediates from about 200 folding and unfolding events (Figure 6 and 7). The results reveal that all four possible hairpin conformations are accessible in the simulations. However, their lifetimes 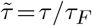 relative to the folded state are slightly different. The hairpin structure formed by the two terminal guanosine repeats (labeled H4 in Figure 6) is most stable 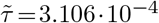.

**Figure 6.**
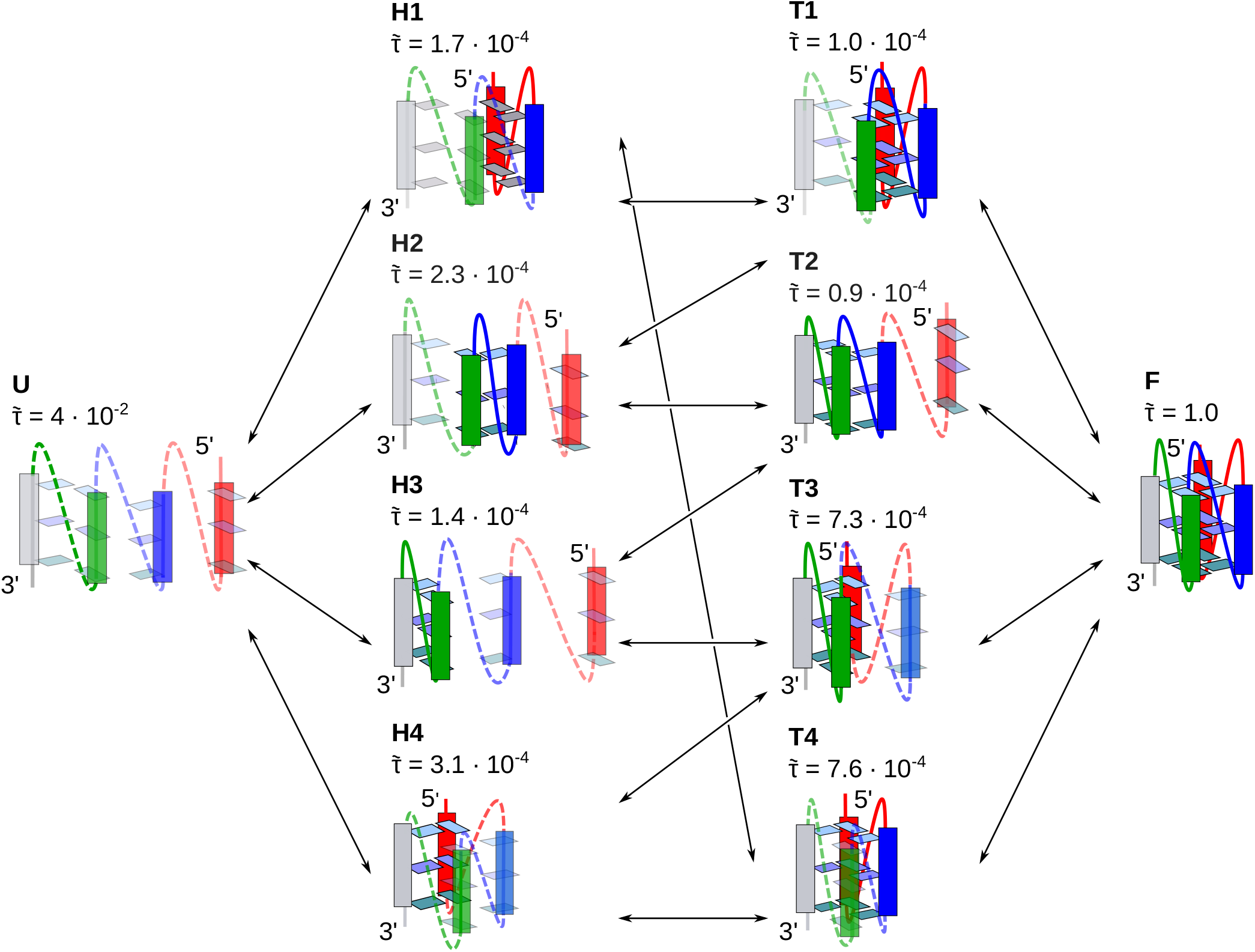
Folding and unfolding pathways of TERRA25. The arrows correspond to the transitions between the conformations. The pathways pass through two intermediate states: hairpin and triplex state. In total four different hairpin (H1, H2, H3, H4) and four different triplex (T1, T2, T3, T4) conformations are observed. The rescaled lifetimes 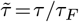 are shown above the structures with *τ*_*F*_ = 6 · 10^−5^ s_CG_. The rescaled lifetimes are related to the experimental timescale via 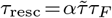 with *α* = 1.56 · 10^7^. The reduced, CG and rescaled lifetimes are listed in the Supporting Information (Table S3).

**Figure 7.**
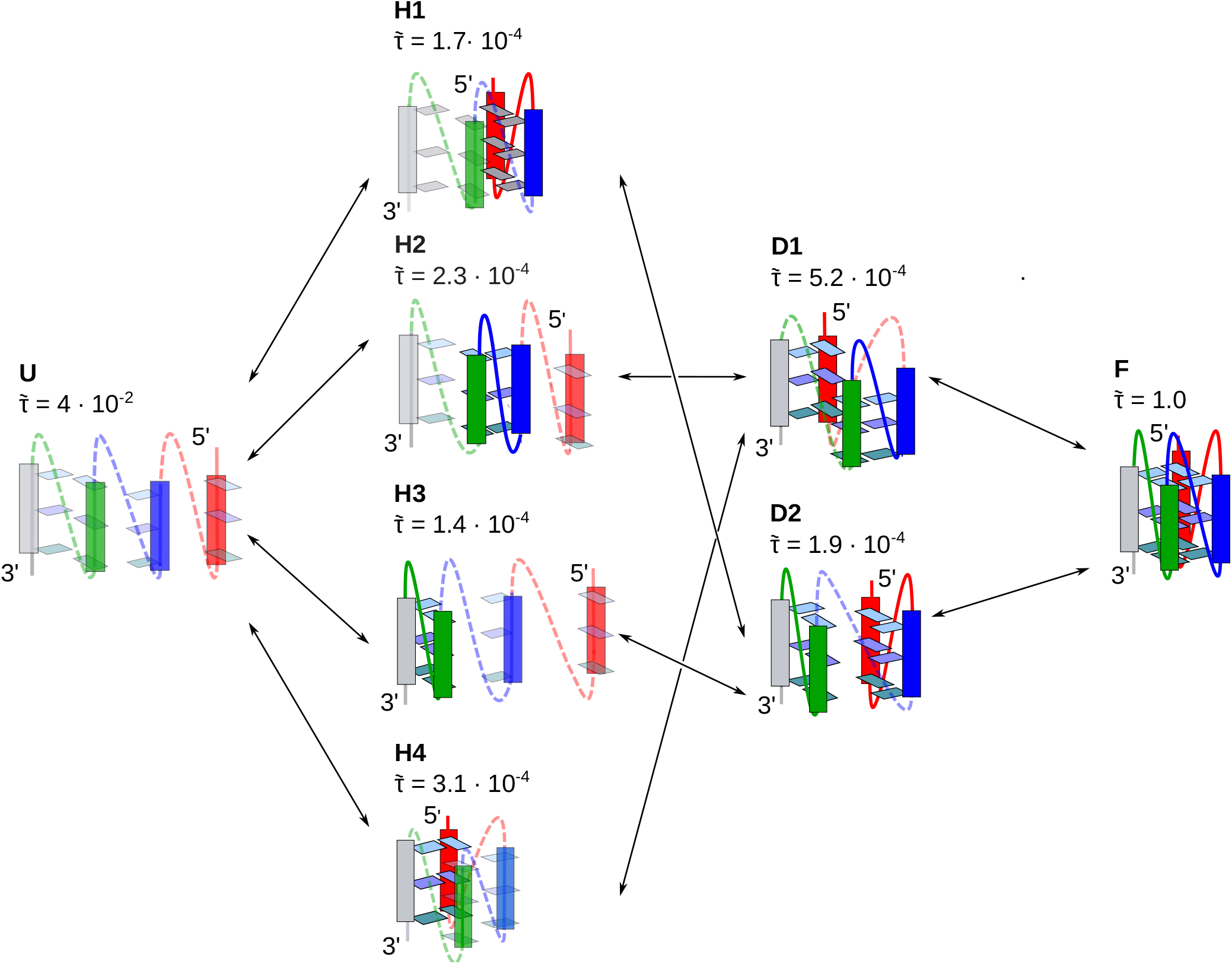
Folding and unfolding pathways of TERRA25. The arrows correspond to the transitions between the conformations. The pathways pass through two intermediate states: hairpin and double-hairpin state. As in Figure 5, four different hairpin (H1, H2, H3, H4) and two different double-hairpin (D1, D2) conformations are observed. The rescaled lifetimes 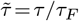 are shown above the structures with *τ*_*F*_ = 6 · 10^−5^ sCG. The rescaled lifetimes are related to the experimental timescale via 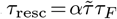 with *α* = 1.56 · 10^7^. The reduced, CG and rescaled lifetimes are listed in the Supporting Information. (Table S3)

Hairpin structures with propeller loops have shorter lifetimes. The shortest lifetime is observed for the hairpin structure formed by the first two repeats at the 3’ end (labeled H3, 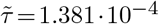).

Moreover, a hairpin intermediate at the 5’ end, similar to H1 in Figure 6 and Figure 7, was recently observed in smFRET experiments on the folding pathway of a plant rG4 (36). The situation is similar for the triplex intermediates. Again, all four possible triplex conformations are observed in the simulations. Triplex intermediates with 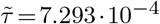; *τ*_resc_ = 0.6986 s (T3) and 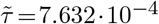 ; *τ*_resc_ = 0.7310 s (T4) are formed by the two terminal repeats and one central repeat (T3 and T4 in Figure 6). The other triplexes (T1, T2) have slightly shorter lifetimes.

Interestingly, a triplex similar to T1 has been observed in smFRET experiments for a plant rG4 (36). Moreover, RNA triplexes with propeller loops have been proposed for rG4s and dG4s with two quartets (35).

In addition to hairpin and triplex, an additional class of intermediates is observed: Two hairpins are created sequentially and form a double-hairpin intermediate state (Figure 7). In pathways containing a double-hairpin intermediate, no triplex state is observed. The double-hairpin structure D1, in which each hairpin comprises one free end, is the most stable 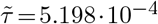; *τ*_resc_ = 0.4980 s .

This is further supported by a similar double-hairpin conformation from NMR spectroscopy on a hybrid G4 of human telomeric DNA (29).

In summary, the pathways between the unfolded and folded state of TERRA25 pass through two intermediate states. The first intermediate is a hairpin and the second one is either a triplex or a double-hairpin state. Each state comprises several different conformations leading to the high conformational entropy of the intermediate states. In total, four different hairpin, four different triplex and two different double-hairpin structures are observed along the folding pathways of TERRA25. Based on the rescaling factor (eq. **3**) the lifetimes of the intermediates are estimated to be on the order of a hundred milliseconds (Table S3). Note however that the rescaling factor provides only a rough estimate due to the coarsening of complex molecular interactions into a single scalar. Still, additional all-atom simulations of the hairpin intermediates reveal that these structures remain stable for more than 500 ns (Figure S7). By contrast, if the dangling ends are removed the unfolding kinetics is much faster in agreement with previous MD simulations (37, 79).

### Conformational entropy as hallmark of rG4 folding

The results for TERRA25 show that the intermediates are degenerate with two to four alternative conformations per state leading to a branched multi-pathway folding process driven by conformational entropy. Here, the immediate question arises whether entropy driven multi-pathway folding is a general feature of rG4 systems. To investigate this hypothesis, we investigate three synthetic rG4 sequences with varying loop length and a plant rG4 sequence with mixed loops (see Table 1). Initially, we calculate the free energy profiles and determine the stability of the five systems. The free energy profiles as function of the radius of gyration *r*_*g*_ all have similar shapes (Figure 8A). The position of the first minimum increases with increasing loop length as expected. The height of the barrier, which separates the folded and unfolded state, increases with decreasing loop length for synthetic G4 sequences indicating that the loop length affects the folding kinetics. The most pronounced differences are the height and width of the second minimum. Clearly, short sequences have a narrower unfolded state due to the smaller number of possible unfolded conformations and hence a smaller conformational entropy. The entropic contribution leads to the highest free energy level for U, followed by UU and UUU. A general trend emerges for the stability namely that rG4 stability decreases with increasing loop length (Figure 8B). The results for UU, UUU and TERRA25 are in quantitative agreement with the experimental results. For the one nucleotide loop deviations are observed which likely result from the different loop sequence used in experiments and simulations.

**Figure 8.**
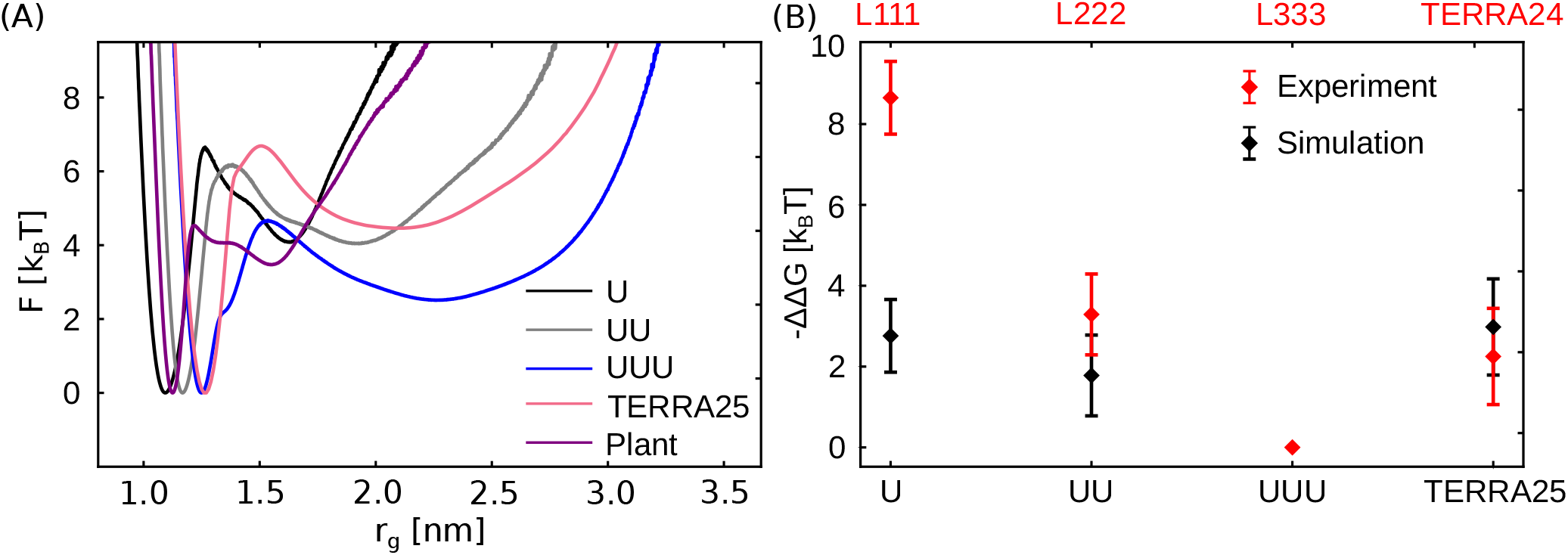
Free energy profiles and stability of different rG4 systems (Table 1). (A) Free energy profile as a function of the radius of gyration *r*_g_ . (B) Gibbs free energy and van’t Hoff Gibbs free energy change for the four different systems. The experimental data was calculated from ∆H-T∆S (see Supporting Information) based on the thermodynamic parameters provided in ref. (74). The Gibbs free energy difference ∆∆*G* =∆*G* −∆*G*ref was calculated with respect to a reference system. For the simulations, we chose the UUU system and the L333 system ref. (74) for the experiments. Errors of results from simulations were calculated from block averaging by dividing trajectories into 10 equal blocks. Errors for experimental values are taken from ref. (74).

The free energy landscapes (Figure 9) for the five systems reveal that the branched multi-pathway folding is a general feature for all rG4 systems investigated here. The order parameters *s*_1_ and *s*_2_ allow us to resolve the different intermediate states and confirm that the 12 intermediate states occur in all rG4 systems. Interestingly, the distribution of the states is not uniform but depends on the exact sequence (see Table S3 in the Supporting Information for the populations in the intermediate states). Again, the H1 intermediate state for the plant rG4 agrees with the predictions from smFRET (36). However, due to the positioning of the labels and the limited resolution in those experiments further work by smFRET is required to resolve all intermediates.

**Figure 9.**
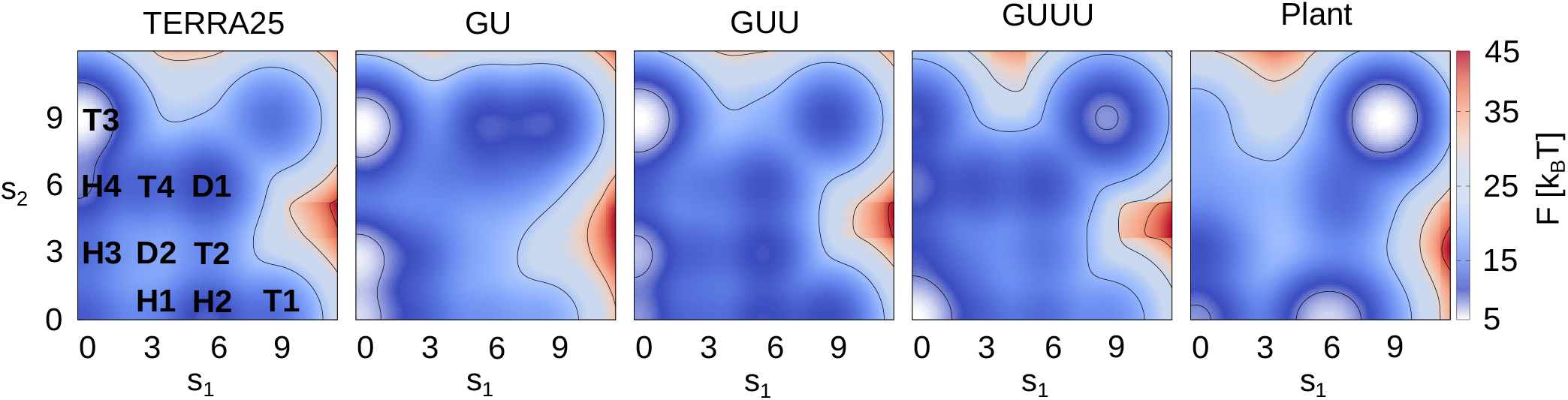
Free energy landscapes of rG4 folding as function of the order parameters *s*_1_ and *s*_2_ (eq. **8**). The positions of the intermediate states are indicated on the left. The free energies of folded and unfolded states are rescaled for visualization.

## CONCLUSION

The folding kinetics of rG4s is essential for their function in cellular processes. In the current work, we resolve the folding pathways of a G4 derived from human telomeric repeat-containing RNA (TERRA25) by combining all-atom MD, CG simulations and CD experiments.

Our results reveal a branched multi-pathway folding process for TERRA25 driven by conformational entropy. Moreover, four rG4 systems with varying loop length highlight the fundamental importance of conformational entropy in rG4 folding and reveal branched multi-pathway folding as a characteristic feature of rG4 systems.

Initially, we developed a matching procedure based on atomistic simulations in explicit water to correctly capture the ionic double layer in the implicit solvent CG simulations. The matching correctly reproduces the local salt concentration in the vicinity of the negatively charged phosphate groups and therefore yields close agreement of the fraction of folded structures obtained from CG simulations and experiments at a given salt concentration.

Folding of rG4s is on the timescale of minutes (21) and therefore out of reach for atomistic simulations. CG simulations using the TIS model, on the other hand, reach beyond this limit and allow us to resolve the folding pathways and intermediate states with sufficient statistics. The simulations reveal on-pathways intermediates corresponding to hairpin, triplex or double-hairpin states for all rG4 systems. The intermediates possess a high conformational entropy with two to four structures contributing to each state. However, properties that are easy to measure such as the end-to-end distance are insufficient to discriminate between the different structures. Here, the insights from the simulations can guide the experiments to detect all intermediate structures, for instance by strategically placed fluorophores in smFRET experiments (35, 36).

In the future, more complex models could be employed to further explore the off-pathway intermediates including strand-shifts or syn-anti G-patterns. In any case, combining atomistic MD, CG simulations and experiments is a valuable starting point to capture the influence of the ionic environment and to resolve the multi-pathway folding landscape.

## SUPPORTING INFORMATION

The Supporting Information is available free of charge. Additional information regarding the methods presented in the main manuscript are included. Additional results regarding non-native intermediates on TERRA25 folding pathway are presented. Tables including lifetimes, populations and transition rates for TERRA25 folding intermediates as well as transition rates, equilibrium constants and Gibbs free energies of five simulated rG4 structures are presented. Additional results regarding all-atom simulations of TERRA25 folding intermediates are presented.

## ACKNOWLEDGMENTS

We acknowledge financial support from the DFG CRC902 and Emmy Noether program (Grant No. 315221747). GOETHE HLR and NHR@FAU are acknowledged for supercomputing access. Work at BMRZ is supported by the state of Hesse. N. S. and M. U. thank Dave Thirumalai, Naoto Hori, Hung T. Nguyen and Debayan Chakraborty for sharing their code and helpful discussions. M. U. thanks Jürgen Köfinger and Sergio Cruz-Leòn for many fruitful discussions.

