## Supporting Information for "RNA G-quadruplex folding is a multi-pathway process driven by conformational entropy"

### RMSD and concentration profiles from MD simulations

We calculated the RMSD for the heavy atoms of TERRA25 with respect to the starting structure. Figure S1A shows the RMSD as a function of time for four selected KCl concentrations (0.48 mM, 0.22 M and 1.03 M) and for 1 M NaCl. The corresponding simulation snapshots after 100 ns are shown in Figure S1 B-E. For high concentrations the simulated structures remain close to the starting structure as expected (RMSD < 3 Å). For very low concentrations, the simulated structures deviate in particular in the loop regions. The concentration profiles for 3 selected KCl concentrations (0.48 mM, 0.22 M, 1.03 M) and 1M NaCl are shown in Fig. S2.

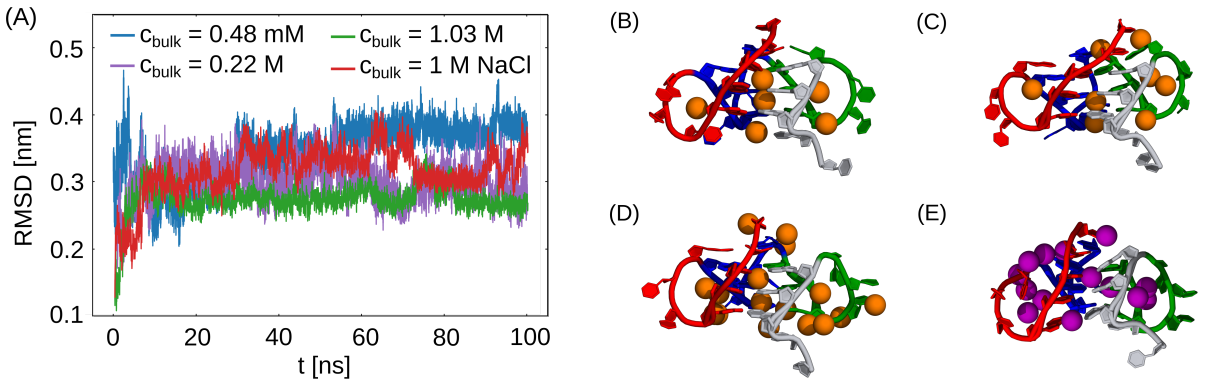


Figure S1: (A) RMSD from all-atom MD simulations of TERRA25 for three selected KCl concentrations, c = 0.48 mM, c = 0.22 M, c = 1.03 M, and c=1 M NaCl. Corresponding snapshots after 100 ns simulation time of TERRA25 in (B) 0.48 mM KCl, (C) 0.22 M KCl, (D) 1.03 M KCl and (E) 1 M NaCl with cations within 0.3 nm from the geometrical center of TERRA25.

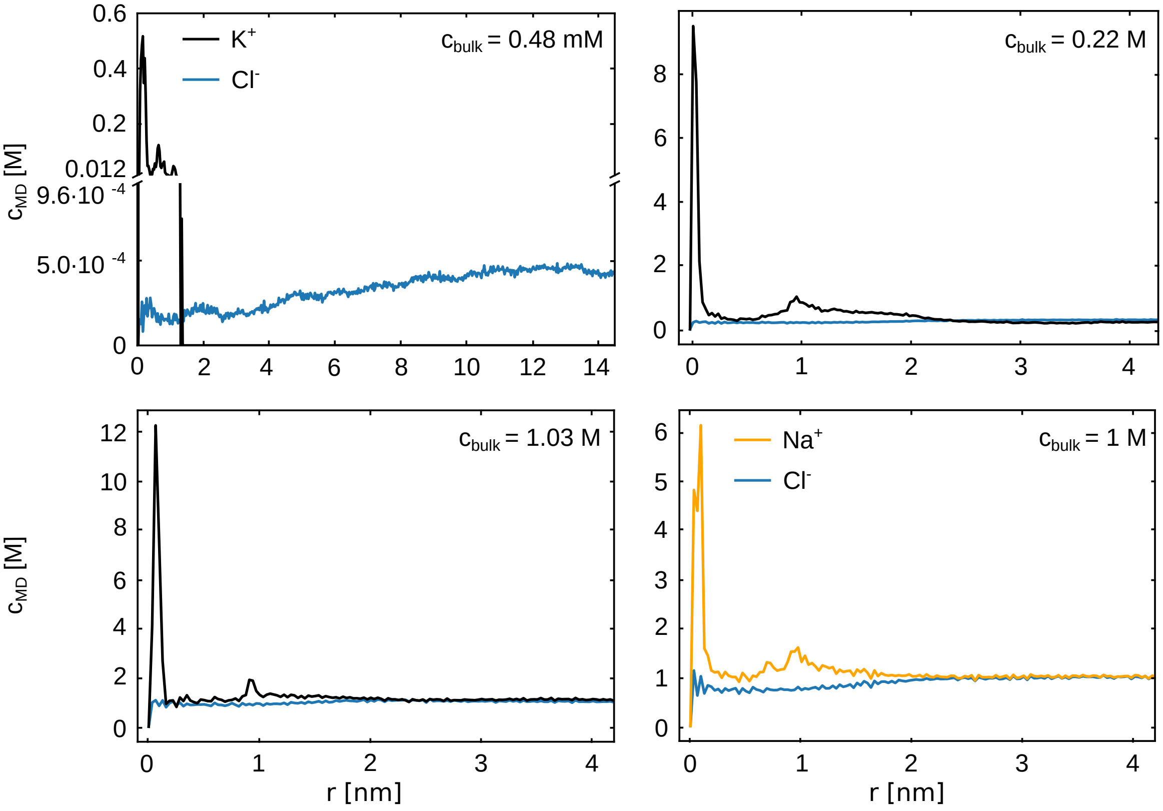


Figure S2: Cation and anion concentration profiles for three selected KCl concentrations and c=1 M NaCl.

Table S1: Values of local concentration $c_{CG}$ and bulk concentration $c_{MD}$ from all-atom simulations of TERRA25 used in linear interpolation.

| $c_{bulk}$ [M] | $c_{CG}$ [M] |
|---|---|
| 0 | 0 |
| 0.00048 | 0.07 |
| 0.016 | 0.18 |
| 0.053 | 0.36 |
| 0.082 | 0.50 |
| 0.14 | 0.66 |
| 0.22 | 0.86 |
| 1.03 | 1.39 |
| 1.89 | 2.33 |

### Non-native intermediates for TERRA25

In addition to hairpin and triplex, intermediates with non-native interactions are frequently observed. These intermediates are helical RNA conformation (Figure S3 A) and strand-shifted triplex conformation (Figure S3 B). We found that these off-pathway intermediates are only transient and unfold quickly, with typical time each state is stable $t_{\text{avg}}$ smaller than 10 ms.

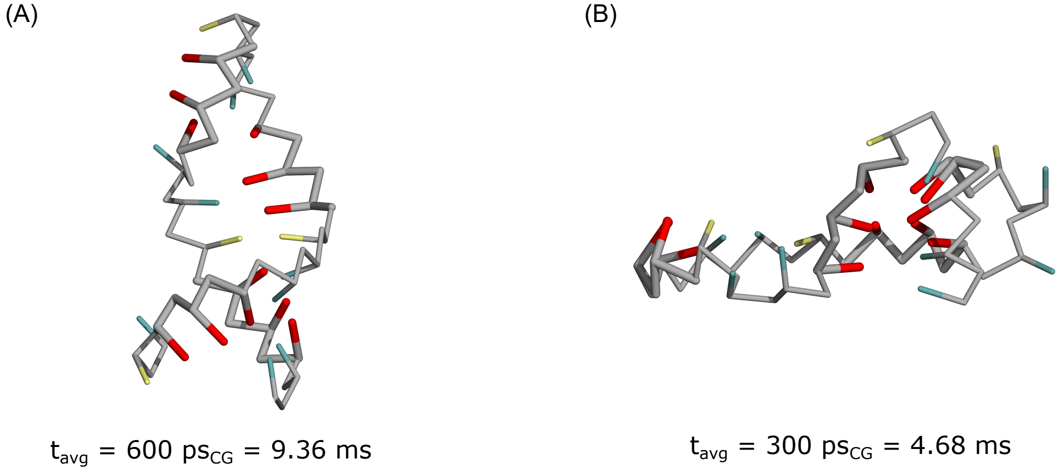


Figure S3: Non-native intermediate conformations for TERRA25. (A) Helical conformation and (B) strand-shifted triplex. The typical time over which the intermediate is stable is given below the structures.

### Rate equations for rG4 folding

The time evolution of the probability of each state is given by

$$\frac{dp_i}{dt} = \sum_{j\neq i} k_{ij} P_j - \sum_{j\neq i} k_{ji} P_i \tag{1}$$

where $p_i$ is the population of state $i$ and $k_{ij}$ are the kinetic rate coefficients. For rG4 folding, we obtain 14 coupled differential equations. The numbers for the rate coefficients and the equilibrium populations are shown in Tables S2 and S3. Note that in equilibrium, the time derivative $dp_i/dt = 0$. With the equations used here, this is the case due to our definition of rates in equation 4 in the main text and populations from equation 3. For example, in the case of the conformation H1, the time evolution of the population is described by equation 2.

$$\begin{aligned}\frac{dp_{\mathrm{H1}}}{dt} &= -k_{\mathrm{T1H1}}\cdot p_{\mathrm{H1}} - k_{\mathrm{D2H1}}\cdot p_{\mathrm{H1}} - k_{\mathrm{UH1}}\cdot p_{\mathrm{H1}} + k_{\mathrm{H1T1}}\cdot p_{\mathrm{T1}} + k_{\mathrm{H1D2}}\cdot p_{\mathrm{D2}} + k_{\mathrm{H1U}}\cdot p_{\mathrm{U}} \\ &= -\frac{N_{\mathrm{T1H1}}}{2t_{\mathrm{H1}}}\cdot\frac{t_{\mathrm{H1}}}{t_{\mathrm{SIM}}} - \frac{N_{\mathrm{D2H1}}}{2t_{\mathrm{H1}}}\cdot\frac{t_{\mathrm{H1}}}{t_{\mathrm{SIM}}} - \frac{N_{\mathrm{UH1}}}{2t_{\mathrm{H1}}}\cdot\frac{t_{\mathrm{H1}}}{t_{\mathrm{SIM}}} + \frac{N_{\mathrm{T1H1}}}{2t_{\mathrm{T1}}}\cdot\frac{t_{\mathrm{T1}}}{t_{\mathrm{SIM}}} \\ &+ \frac{N_{\mathrm{D2H1}}}{2t_{\mathrm{D2}}}\cdot\frac{t_{\mathrm{D2}}}{t_{\mathrm{SIM}}} + \frac{N_{\mathrm{UH1}}}{2t_{\mathrm{U}}}\cdot\frac{t_{\mathrm{U}}}{t_{\mathrm{SIM}}}\end{aligned} \tag{2}$$

It is clear from equation that the positive and negative parts cancel out and the result is zero. The same is true for other intermediate conformations.

Table S2: Rates of transitions between intermediates on the TERRA25 folding pathway. Each transition is labeled by the name of intermediates. Therefore, $k_{ij}$ is the rate of transition from state $j$ to state $i$. CG rates can be related to the experimental rates by using the scaling factor $\alpha = 1.56 \cdot 10^7$ (see equation 3 in the main text). The experimental folding rate for TERRA25 is $k_{\mathrm{FU}} = 2.4 \cdot 10^{-2}$.[S1] Errors were calculated from block averaging where of all trajectories were divided into 10 equal blocks.

| State $i$ | State $j$ | $k_{ij}$ [ $10^7$ $\mathrm{s}_{\mathrm{CG}}^{-1}$] | $k_{ji}$ [ $10^7$ $\mathrm{s}_{\mathrm{CG}}^{-1}$ ] | rescaled $k_{ij}$ [ $\mathrm{s}^{-1}$] | rescaled $k_{ji}$ [ $\mathrm{s}^{-1}$] |
|---|---|---|---|---|---|
| H1 | U | 0.0017 ± 0.0005 | 6.0 ± 2.0 | 0.0011 ± 0.0003 | 4.0 ± 1.0 |
| H2 | U | 0.0013 ± 0.0004 | 4.0 ± 1.0 | 0.008 ± 0.002 | 2.7 ± 0.8 |
| H3 | U | 0.007 ± 0.002 | 6.0 ± 2.0 | 0.005 ± 0.001 | 4.0 ± 1.0 |
| H4 | U | 0.0012 ± 0.0004 | 2.3 ± 0.8 | 0.008 ± 0.002 | 1.8 ± 0.5 |
| T1 | H1 | 3.0 ± 1.0 | 1.0 ± 0.3 | 2.3 ± 0.7 | 0.7 ± 0.2 |
| T1 | H2 | 2.3 ± 0.7 | 7.0 ± 2.0 | 1.6 ± 0.5 | 4.0 ± 1.0 |
| T2 | H2 | 0.6 ± 0.2 | 5.0 ± 2.0 | 0.4 ± 0.1 | 4.0 ± 1.0 |
| T2 | H3 | 1.2 ± 0.4 | 4.0 ± 1.0 | 0.8 ± 0.2 | 2.7 ± 0.8 |
| T3 | H3 | 5.0 ± 1.0 | 0.4 ± 0.1 | 3.0 ± 0.9 | 2.8 ± 0.8 |
| T3 | H4 | 1.7 ± 0.5 | 0.6 ± 0.2 | 1.1 ± 0.3 | 0.4 ± 0.1 |
| T4 | H4 | 0.29 ± 0.09 | 1.2 ± 0.4 | 0.19 ± 0.06 | 0.8 ± 0.3 |
| F | T1 | 8.0 ± 2.0 | 0.0004 ± 0.0001 | 5.0 ± 2.0 | 0.00028 ± 0.00008 |
| F | T2 | 9.0 ± 3.0 | 0.00017 ± 0.00005 | 6.0 ± 2.0 | 0.00012 ± 0.00003 |
| F | T3 | 1.2 ± 0.3 | 0.0008 ± 0.0002 | 0.8 ± 0.2 | 0.0005 ± 0.0002 |
| F | T4 | 1.2 ± 0.4 | 0.00007 ± 0.00002 | 0.8 ± 0.2 | 0.0004 ± 0.0001 |
| D1 | H4 | 0.7 ± 0.2 | 1.3 ± 0.4 | 0.5 ± 0.1 | 0.9 ± 0.3 |
| D2 | H1 | 1.1 ± 0.3 | 2.9 ± 0.9 | 0.8 ± 0.2 | 1.9 ± 0.6 |
| D2 | H3 | 1.4 ± 0.4 | 1.4 ± 0.4 | 0.9 ± 0.3 | 1.0 ± 0.3 |
| F | D1 | 1.8 ± 0.5 | 0.00022 ± 0.00007 | 1.2 ± 0.4 | 0.00015 ± 0.00004 |
| F | D2 | 4.0 ± 1.0 | 0.000025 ± 0.000007 | 2.9 ± 0.8 | 0.000017 ± 0.000005 |
| F | U | 0.0375 ± 0.0164 | 0.00163 ± 0.00060 | 0.0240 ± 0.0105 | 0.00105 ± 0.00039 |

### Populations and lifetimes of the states

We calculate population $P_i$ of state $i$ by dividing time spent in the state i $t_i$ by the total simulation time $t_{\text{sim}}$

$$P_i = \frac{t_i}{t_{\text{sim}}} \tag{3}$$

The lifetimes are calculated according to equation 5 in the main manuscript. All values are listed in Table S3.

Table S3: Populations and lifetimes of the folded, unfolded and all intermediate states obtained from the CG simulations. For the reduced lifetimes the lifetime of the folded state ($\tau = 60\ \mu\text{s}_{\text{CG}}$) was used as reference. The lifetime in CG time units can be related to the experimental timescale via the scaling factor $\alpha = 1.56 \cdot 10^7$ (see equation 3 in the main text). Errors for results from simulations were calculated from block averaging where of all trajectories were divided into 10 equal blocks.

| State | population | | | reduced lifetime $\tilde{\tau}$ | | | lifetime $\tau$ [$\text{s}_{\text{CG}}$] | | | rescaled $\tau$ [s] | | |
|---|---|---|---|---|---|---|---|---|---|---|---|---|
| F | $9.51 \cdot 10^{-1}$ | ± | $2.1 \cdot 10^{-3}$ | 1.0 | ± | 0.1 | $6.14 \cdot 10^{-5}$ | ± | $6.1 \cdot 10^{-6}$ | 957.8 | ± | 95.1 |
| U | $4.81 \cdot 10^{-2}$ | ± | $2.8 \cdot 10^{-4}$ | 0.04 | ± | 0.01 | $2.7 \cdot 10^{-6}$ | ± | $6.4 \cdot 10^{-7}$ | 42.0 | ± | 9.6 |
| H1 | $1.39 \cdot 10^{-5}$ | ± | $3.2 \cdot 10^{-7}$ | $1.686 \cdot 10^{-4}$ | ± | $1.7 \cdot 10^{-7}$ | $10.35 \cdot 10^{-9}$ | ± | $1.1 \cdot 10^{-11}$ | 0.1615 | ± | $1.7 \cdot 10^{-4}$ |
| H2 | $1.48 \cdot 10^{-4}$ | ± | $2.4 \cdot 10^{-6}$ | $2.318 \cdot 10^{-4}$ | ± | $2.0 \cdot 10^{-7}$ | $14.23 \cdot 10^{-9}$ | ± | $1.2 \cdot 10^{-11}$ | 0.2220 | ± | $1.9 \cdot 10^{-4}$ |
| H3 | $5.75 \cdot 10^{-5}$ | ± | $8.4 \cdot 10^{-7}$ | $1.381 \cdot 10^{-4}$ | ± | $2.9 \cdot 10^{-7}$ | $8.48 \cdot 10^{-9}$ | ± | $1.8 \cdot 10^{-11}$ | 0.1323 | ± | $2.8 \cdot 10^{-4}$ |
| H4 | $2.21 \cdot 10^{-4}$ | ± | $3.0 \cdot 10^{-6}$ | $3.106 \cdot 10^{-4}$ | ± | $3.4 \cdot 10^{-7}$ | $19.07 \cdot 10^{-9}$ | ± | $2.1 \cdot 10^{-11}$ | 0.2975 | ± | $3.3 \cdot 10^{-4}$ |
| T1 | $4.86 \cdot 10^{-5}$ | ± | $9.5 \cdot 10^{-7}$ | $1.001 \cdot 10^{-4}$ | ± | $2.3 \cdot 10^{-7}$ | $6.15 \cdot 10^{-9}$ | ± | $1.4 \cdot 10^{-11}$ | 0.0960 | ± | $2.2 \cdot 10^{-4}$ |
| T2 | $1.76 \cdot 10^{-5}$ | ± | $4.3 \cdot 10^{-7}$ | $0.868 \cdot 10^{-4}$ | ± | $2.0 \cdot 10^{-7}$ | $5.33 \cdot 10^{-9}$ | ± | $1.2 \cdot 10^{-11}$ | 0.0832 | ± | $1.9 \cdot 10^{-4}$ |
| T3 | $6.1 \cdot 10^{-4}$ | ± | $1.1 \cdot 10^{-5}$ | $7.293 \cdot 10^{-4}$ | ± | $2.8 \cdot 10^{-7}$ | $44.78 \cdot 10^{-9}$ | ± | $1.7 \cdot 10^{-11}$ | 0.6986 | ± | $2.7 \cdot 10^{-4}$ |
| T4 | $5.17 \cdot 10^{-5}$ | ± | $1.3 \cdot 10^{-6}$ | $7.632 \cdot 10^{-4}$ | ± | $3.8 \cdot 10^{-7}$ | $46.86 \cdot 10^{-9}$ | ± | $2.3 \cdot 10^{-11}$ | 0.7310 | ± | $3.6 \cdot 10^{-4}$ |
| D1 | $1.16 \cdot 10^{-4}$ | ± | $6.1 \cdot 10^{-6}$ | $5.198 \cdot 10^{-4}$ | ± | $5.1 \cdot 10^{-7}$ | $31.92 \cdot 10^{-9}$ | ± | $3.1 \cdot 10^{-11}$ | 0.4980 | ± | $4.8 \cdot 10^{-4}$ |
| D2 | $5.5 \cdot 10^{-4}$ | ± | $1.3 \cdot 10^{-6}$ | $1.894 \cdot 10^{-4}$ | ± | $1.07 \cdot 10^{-7}$ | $11.630 \cdot 10^{-9}$ | ± | $6.6 \cdot 10^{-12}$ | 0.1814 | ± | $1.0 \cdot 10^{-4}$ |

### Gibbs free energy from CG simulations and thermal melding curves

The Gibbs free energy is related to the affinity constant via

$$\Delta G = -k_B T \ln(K_{\text{eq}}) \tag{4}$$

where the equilibrium constant $K_{eq}$ is calculated from the rate constants using $K_{eq} = k_{FU}/k_{UF}$. The values are listed in Table S4 for the 5 different systems.

From UV melting experiments,[S2] the Gibbs free energy can be extracted from the van't Hoff representation $\ln(K_{eq})$ vs 1/T. Note that the approach is only valid for two-state dynamics and reversible reactions. In ref.[S2] the van't Hoff enthalpy $\Delta H_{VH}$, entropy $\Delta S_{VH}$ and Gibbs free energy $\Delta G_{VH}$ at 37 °C and for 20 $\mu$M RNA in 5 mM KCl were determined for the synthetic rG4s investigated here.

Since our simulations were performed at 298.15 K, we calculate the Gibbs free energy 298.15 K from the entropy change $\Delta S_{VH}$ and enthalpy change $\Delta H_{VH}$. Since the volume remains constant in the CG folding simulations, the simulation results obtained from equation 4 can be compared directly to $\Delta H_{VH} - T\Delta S_{VH}$. The values are listed in Table S4. Simulations were done at 0.3 mM KCl bulk concentration for 10 $\mu$M RNA concentration.

Table S4: Thermodynamic and kinetic properties for the rG4 systems. $k_{FU}$, $k_{UF}$, $K_{eq}$ and $\Delta G$ from simulations. The values for the Gibbs free energy $\Delta H_{VH} - T\Delta S_{VH}$ are calculated from the experimental values in ref.[S2] for 298.15 K. The experiments were done at 20 $\mu$M RNA in 5 mM potassium chloride. Note that the experiments are for rG4 libraries containing mixed loop sequences with a defined length, and not only uridine nucleosides. Simulations were done at 0.3 mM KCl bulk concentration for 10 $\mu$M RNA concentration. The plant sequence was simulated at 0.08 M KCl. Errors for results from simulations were calculated from block averaging where of all trajectories were divided into 10 equal blocks.

| rG4 | $k_{FU}$ [ $10^5$ $s_{CG}^{-1}$] | $k_{UF}$ [ $10^4$ $s_{CG}^{-1}$ ] | $K_{eq}$ | $\Delta G[k_BT]$ | $k_{FU}^{exp}$ $[s^{-1}]$ | rG4 library | $\Delta H_{VH} - T\Delta S_{VH}$ $[k_BT]$ |
|---|---|---|---|---|---|---|---|
| U | 53.09 ± 6.72 | 28.74 ± 3.89 | 18.47 ± 0.41 | -2.92 ± 0.19 | | L111 | -17.21 ± 0.7 |
| UU | 10.91 ± 3.24 | 15.68 ± 2.92 | 6.96 ± 2.61 | -1.94 ± 0.29 | | L222 | -11.85 ± 0.6 |
| UUU | 1.50 ± 0.31 | 12.77 ± 14.68 | 1.17 ± 0.82 | -0.16 ± 0.71 | | L333 | -8.56 ± 0.2 |
| TERRA25 | 3.75 ± 1.64 | 1.63 ± 0.60 | 23.5 ± 13.65 | -3.14 ± 0.475 | $-2.4 \cdot 10^{-2}$ | TERRA | -10.81 ± 0.1 |
| Plant | 15.10 ± 3.19 | 21.94 ± 4.45 | 6.88 ± 2.35 | -1.93 ± 0.36 | | | |

### Order parameters

We defined repeat order parameter $r_{ij}$ and two additional order parameters $s_1$ and $s_2$ to discriminate between different intermediates (see Methods in the main manuscript). Figure S4 represents values of the order parameters in the intermediate states.

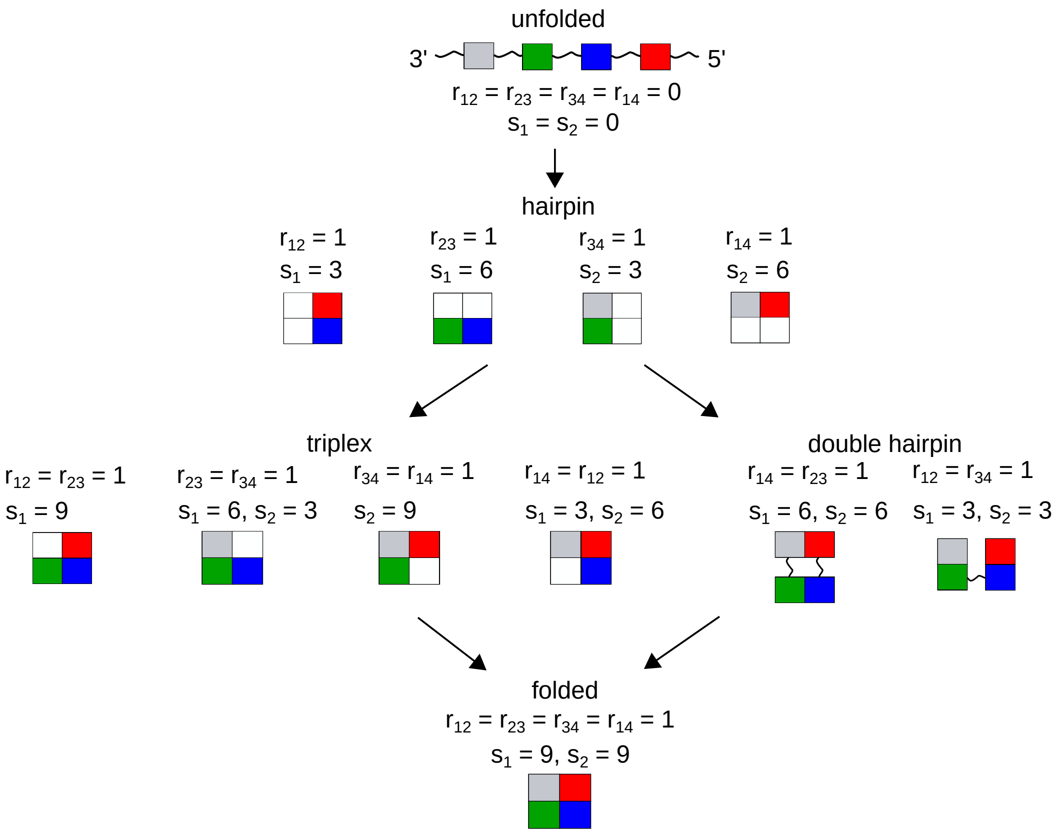


Figure S4: Scheme representing repeat order parameters $r_{\mathrm{ij}}$ and intermediate-defining order parameters $s_1$ and $s_2$. Guanosine repeats are defined as cubes and the folded repeats are indicated in color. Only non-zero order parameters are indicated for intermediates.

### Native polyacrylamide gel electrophoreses (PAGE)

Correct folding of TERRA25 was characterized via native PAGE (15% acrylamide gel). 400 pmol were loaded in 50% glycerol and 5 mM KCl. Gels were prepared in 50 mM Tris-borate buffer (pH = 8.4) supplemented with 5 mM KCl. Bands were separated in the same buffer at 0.8 W for 90 min with water cooling (13 °C water temperature). The bands have been visualized by Stains-All.

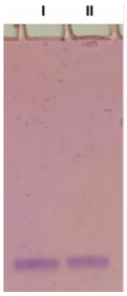


Figure S5: Native polyacrylamid gel electrophoreses at 13 °C with samples of TERRA25 before (I) and after (II) melting experiments. TERRA25 shows one distinct band and was intact throughout the measurements.

### Melting and annealing in CD melting experiments

Figure S6 shows melting and annealing curves for c=144 mM KCl. The shift of the annealing curve with respect to the melting curve is small, confirming that folding and unfolding proceed along the same pathway. We applied a temperature ramp of 0.5 °C/min. The reason for the small difference in the annealing and melting, an effect known as hysteresis, may imply folding barriers when going from the higher temperature to the lower temperature.

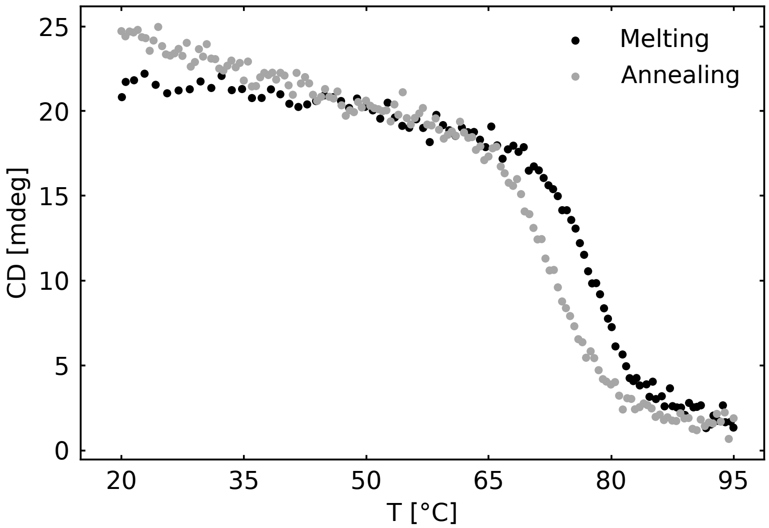


Figure S6: CD melting and annealing curves at 144 mM KCl.

### All-atom simulations of TERRA25 intermediates

#### Modeling of the intermediates

We modeled TERRA25 folding intermediate conformations H1 and H2 as well as their minimum models $H1_{MM}$ and $H2_{MM}$. We created minimum models of H1 and H2 by taking only the appropriate hairpins from the TERRA25 structure. Therefore, the minimum model of H1 ($H1_{MM}$) has the sequence: UAGGGUUAGGG and the minimum model of H2 ($H2_{MM}$) has the sequence: GGGUUAGGG. Additionally, we created full models by using the ModeRNA homology modeling server with the minimum models as a reference. Therefore, we created the full model of H1 with hairpin formed by the two guanosine repeats at the 5' end, while the rest of the sequence was unfolded. Similarly, the H2 full structure consisted of hairpin formed by the two middle guanosine repeats while the rest of the structure was unfolded. In total, four models were created.

#### All-atom MD simulations

The MD simulations were performed with Gromacs version 2020.6 and 2023.1.[S3] Periodic boundary conditions were used with the particle mesh Ewald method[S4] to calculate long range electrostatics, with cubic interpolation and a Fourier spacing of 0.12 nm. The cutoff for Coulomb and Lennard-Jones interactions was 1.2 nm and long-range dispersion corrections for energy and pressure were applied. Bonds to hydrogens were constrained with the LINCS algorithm[S5] with order of four for the matrix inversion correction and one iterative correction. 2 fs time step was used. We used parameters from Amber99sb - ildn* force field[S6] with parambsc0[S7] and $\chi_{OL3}$[S8] corrections and TIP3P water model.[S9] Optimized parameters were used for $K^+$ and $Cl^-$ ions.[S10]

Each model was placed in a cubic box. The dimensions of boxes were chosen with 2.0 nm distance to the edge of the box from the RNA. Therefore, in the $H1_{MM}$ simulation, the box edge length was 7 nm, with 10871 water molecules. The box edge length for the $H2_{MM}$

simulation was 6.6 nm and the box contained 9199 water molecules. For the full hairpin model H1, the box edge length was 10.8 nm and the box contained 40396 water molecules. Finally, the box edge length in the full model simulation of H2 was 9.6 nm and the box contained 29158 water molecules. 25 $K^+$ and 1 $Cl^-$ ion were added to the box. Hence the systems are neutral, and the low concentration promotes unfolding.

Each system was equilibrated in NVT and NPT simulations. The NVT equilibration was done in two parts. During the first 1 ns restraints of 2000 kJ/(mol nm$^2$) were applied to ions and restraints of 1000 kJ/(mol nm$^2$) were applied on the heavy atoms of the hairpins. In the subsequent 1 ns, the restraints on the ions and the hairpins were released. As thermostat the velocity rescaling algorithm with a stochastic term[S11] and with a coupling constant of 0.1 ps was used. During the 2 ns NPT equilibration, pressure coupling was performed using the Berendsen barostat[S12] with a 1 ps coupling constant. In the production runs, the velocity rescaling thermostat[S11] and the Parrinello-Rahman barostat[S13] with a 2 ps coupling constant were used. Each production run was 500 ns long.

The hydrogen bond order parameter $N_{HB}$ was calculated to count the Hoogsteen hydrogen bonds between the hairpin-forming guanosines in each of the models. The coordination function of PLUMED[S14] was used to count the hydrogen bonds. Each pair of guanosines consists of two hydrogen bonds. In total, six hydrogen bonds are formed during hairpin formation.

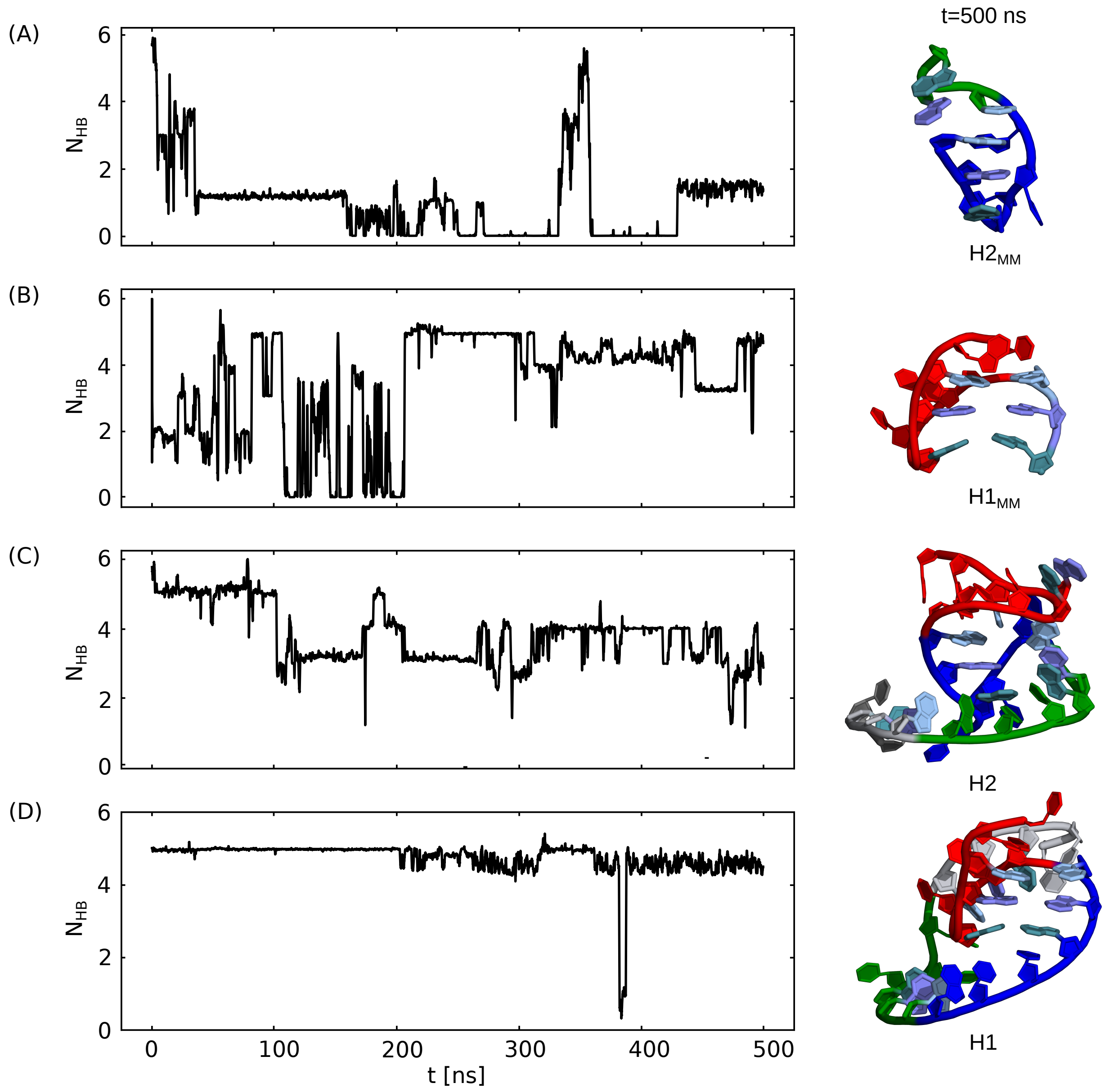


Figure S7: Left panel shows results for number of hydrogen bonds between hairpin-forming guanosines $N_{HB}$ from all-atom simulations of the four hairpin models: $H2_{MM}$ (A), $H1_{MM}$ (B), H2 (C) and H1 (D). Snapshots of structures after 500 ns simulation time are shown on the right.

### References


(S1) Müller, D.; Bessi, I.; Richter, C.; Schwalbe, H. The folding landscapes of human telomeric RNA and DNA G-quadruplexes are markedly different. *Angew Chem. Int.*

*Ed.* **2021**, *60*, 10895–10901.

(S2) Zhang, A. Y.; Bugaut, A.; Balasubramanian, S. A sequence-independent analysis of the loop length dependence of intramolecular RNA G-quadruplex stability and topology. *Biochemistry* **2011**, *50*, 7251–7258.

(S3) Hess, B.; Kutzner, C.; van der Spoel, D.; Lindahl, E. GROMACS 4: Algorithms for Highly Efficient, Load-Balanced, and Scalable Molecular Simulation. *J. Chem. Theo. Comp.* **2008**, *4*, 435–447.

(S4) Darden, T.; York, D.; Pedersen, L. Particle mesh Ewald: An N·log(N) method for Ewald sums in large systems. *J. Chem. Phys.* **1993**, *98*, 10089–10092.

(S5) Hess, B.; Bekker, H.; Berendsen, H. J.; Fraaije, J. G. LINCS: A Linear Constraint Solver for molecular simulations. *J. Comput. Chem.* **1997**, *18*, 1463–1472.

(S6) Bayly, C. I.; Merz, K. M.; Ferguson, D. M.; Cornell, W. D.; Fox, T.; Caldwell, J. W.; Kollman, P. A.; Cieplak, P.; Gould, I. R.; Spellmeyer, D. C. A Second Generation Force Field for the Simulation of Proteins, Nucleic Acids, and Organic Molecules. *J. Am. Chem. Soc.* **1995**, *117*, 5179–5197.

(S7) Pérez, A.; Marchán, I.; Svozil, D.; Sponer, J.; Cheatham, T. E.; Laughton, C. A.; Orozco, M. Refinement of the AMBER force field for nucleic acids: Improving the description of $\alpha/\gamma$ conformers. *Biophys. J.* **2007**, *92*, 3817–3829.

(S8) Zgarbová, M.; Otyepka, M.; Šponer, J.; Mládek, A.; Banáš, P.; Cheatham, T. E.; Jurečka, P. Refinement of the Cornell et al. Nucleic acids force field based on reference quantum chemical calculations of glycosidic torsion profiles. *J. Chem. Theory Comput.* **2011**, *7*, 2886–2902.

(S9) Jorgensen, W. L.; Chandrasekhar, J.; Madura, J. D.; Impey, R. W.; Klein, M. L.

Comparison of simple potential functions for simulating liquid water. *The J. Chem. Phys.* **1983**, *79*, 926–935.

(S10) Mamatkulov, S.; Schwierz, N. Force fields for monovalent and divalent metal cations in TIP3P water based on thermodynamic and kinetic properties. *J. Chem. Phys.* **2018**, *148*.

(S11) Bussi, G.; Donadio, D.; Parrinello, M. Canonical sampling through velocity rescaling. *J. Chem. Phys.* **2007**, *126*, 14101.

(S12) Berendsen, H. J.; Postma, J. P.; Van Gunsteren, W. F.; Dinola, A.; Haak, J. R. Molecular dynamics with coupling to an external bath. *J. Chem. Phys.* **1984**, *81*, 3684–3690.

(S13) Parrinello, M.; Rahman, A. Polymorphic transitions in single crystals: A new molecular dynamics method. *J. Appl. Phys.* **1981**, *52*, 7182–7190.

(S14) Abrams, J. B.; Tuckerman, M. E. Dynamics without Coordinate Transformations. *J. Phys. Chem. B* **2008**, *112*, 15742–15757.
